# Characterisation of O-acetylserine sulfhyrdrylase (CysK) enzymes from bacteria lacking a sulfate reduction pathway

**DOI:** 10.1101/2025.03.26.645513

**Authors:** Jack McGarvie, Keely Oldham, Annmaree Warrender, Erica Prentice, Joanna L. Hicks

## Abstract

Sulfur metabolism plays an important role in bacterial pathogenesis. Elucidation of differences in sulfur metabolism across bacterial pathogens furthers our understanding of host survival and offers opportunities to disrupt these pathways for new therapies. Withing bacteria sulfur metabolism converges at the synthesis of L-cysteine. One of the key mechanisms of obtaining sulfur for the synthesis of L-cysteine is the successive reduction of sulfate to sulfide via the sulfate reduction pathway. Accordingly, L-cysteine biosynthesis is a critical metabolic pathway for bacterial survival, particularly in pathogenic species such as *Neisseria gonorrhoeae* and *Staphylococcus aureus*, which lack the sulfate reduction pathway. O-acetylserine sulfhydrylase catalyses the second step of the two-step synthesis reaction, condensing sulfide or thiosulfate (in the case of OASS-A/CysK or OASS-B/CysM respectively) with O-acetylserine to synthesize cysteine. Here we investigate the enzymatic properties and functional characterization of O-acetylserine sulfhydrylase, from *N. gonorrhoeae* and *S. aureus*, with a focus on substrate specificity, kinetic parameters, and cysteine synthase complex (CSC) formation. Using small angle X-ray scattering and kinetic assays we demonstrate that both *N. gonorrhoeae* and *S. aureus* CysK enzymes utilise only sodium sulfide for the synthesis of cysteine, despite the lack of a sulfate reduction pathway (to generate sulfide) in these organisms. Both enzymes demonstrate a higher affinity for *O*-acetylserine (OAS) compared to sodium sulfide (Na_2_S). We also show that the two cysteine synthesis enzymes, CysE and CysK that traditionally form the cysteine synthase complex do not form a complex in *N. gonorrhoeae*. These findings highlight the functional divergence in sulfur metabolism strategies among bacteria lacking sulfate reduction and provide deeper insights into the adaptive mechanisms of *N. gonorrhoeae* and *S. aureus* in sulfur flux.

## Introduction

Sulfur, biblically referred to as brimstone, is the fifth most common element found on earth [1]. Sulfur is readily available in many different forms, including the inorganic compounds elemental sulfur (S_8_), sulfate (SO_4_^2−^), sulfide (S^2−^), sulfite (SO_3_^2−^), thiosulfate (S O ^2−^), and the polythionates (S O ^2−^; S O ^2−^) [1]. Sulfur also plays a vital role within central biochemistry as a redox active and structural element, due to its wide array of stable oxidation states [1, 2]. The survival of bacteria is dependent on sulfur metabolism with sulfur comprising 0.5-1% of microbial cells dry weight, found in amino acids (e.g. methionine and cysteine), enzyme co-factors (e.g. coenzyme A/M, lipoic acid, biotin, and thiamine), and the foundations of redox reactions (e.g. iron-sulfur complexes, and the redox-active components of disulfide bonds) [2, 3].

Bacterial sulfur metabolism is crucial for bacterial survival. Import of sulfur containing metabolites for biosynthetic processes is derived from assimilation of inorganic sulfate by plants and bacteria [2]. *De novo* biosynthesis of L-cysteine is the main pathway for bacteria to acquire environmental inorganic sulfur. Within most bacteria, cysteine biosynthesis begins with the import of sulfate into the cell where it is subsequently reduced via the reductive sulfate assimilation pathway (RSAP) [4]. The major facilitator superfamily (MFS) and the ATP-binding cassette (ABC) superfamily are the two main families of sulfate transporter proteins responsible for the uptake of inorganic sulfate into microbial cells, via active import [5, 6]. Once inside the cell, successive reduction of sulfate prepares the sulfide for incorporation into L-cysteine (Figure 1). *Escherichia coli* first reduces sulfate to adenosine 5’-phosphosulfate (APS) using the multi-enzyme complex ATP sulfyhdrylase (CysDN) [7]. In most bacteria, APS is reduced to phosphoadenosylphosphosulfate (PAPS) by ATP-kinase (CysC), and PAPS reductase (CysH) subsequently reduces PAPS to sulfite and phosphoadenosylphosphate (PAP) [7, 8]. In *Neisseria* species, the reduction of APS to PAPS, and subsequent reduction to sulfite and PAP, is completed in a single step reaction by APS reductase[9]. Finally, sulfite reductase (CysIJ) reduces sulfite to sulfide [8] which is subsequently combined with *O*-acetylserine, forming L-cysteine (Figure 1).

**Figure 1:**
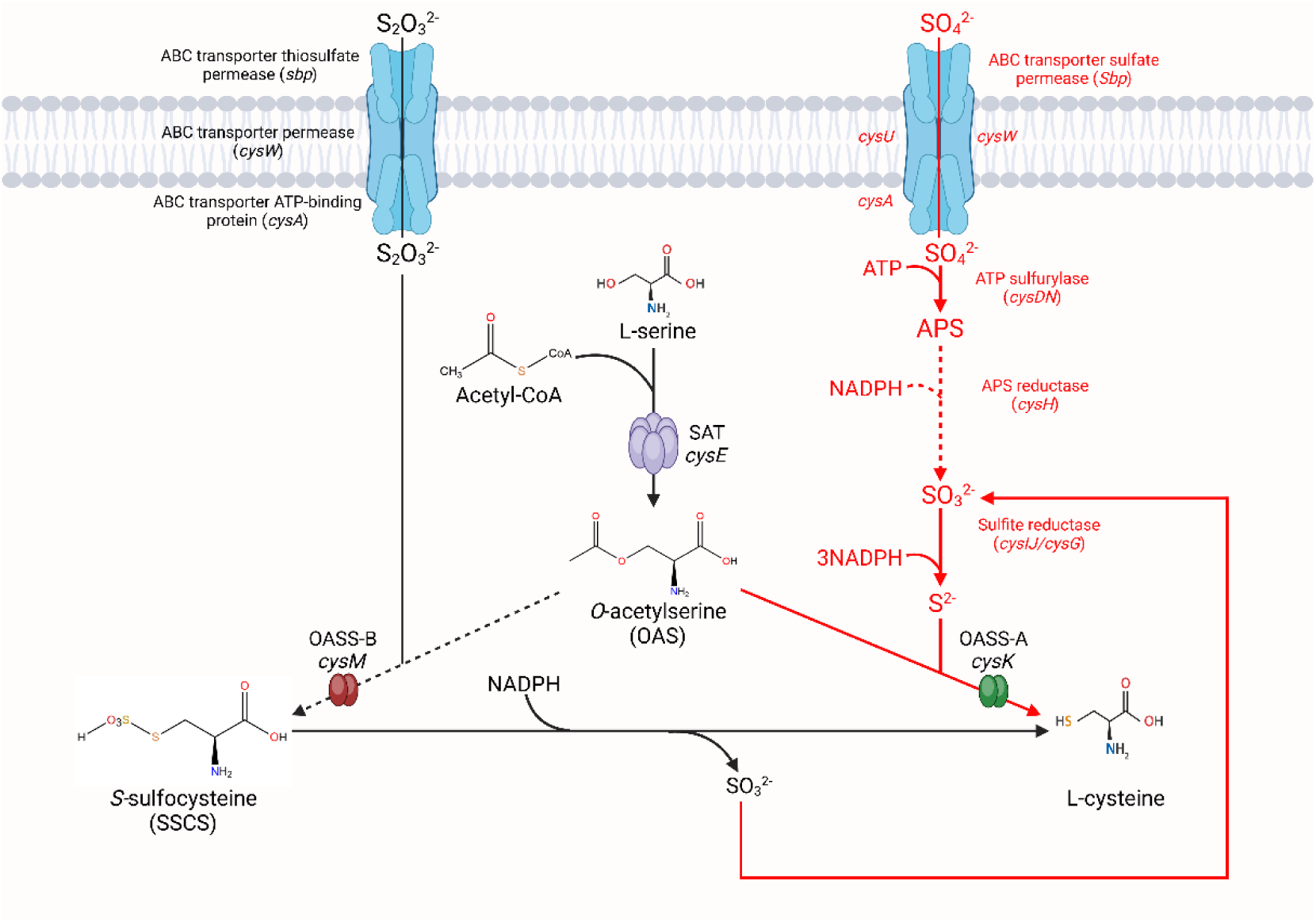
Sulfate acquisition and cysteine biosynthetic pathways in *Neisseria* species

Interestingly, some bacterial pathogens are missing sections of the sulfate reduction pathway, including the notable pathogens *Neisseria gonorrhoeae* and *Staphylococcus aureus*. *N. gonorrhoeae* has only a single sulfate uptake system from the ABC transporter superfamily in its genome [10] composed of a transporter permease (*cysW*), an ATP-binding protein (*cysA*), a periplasmic sulfate-binding protein (*sbp*), and a sulfate permease (*cysU*) which is present in all *N. gonorrhoeae* strains except for FA1090 [6, 10]. However, there is a 3.5 kb deletion between the *cysG* and *cysN* genes in all *N. gonorrhoeae* strains, resulting in the removal of *cysH* and *cysD* from this operon [10]. These genes are not present anywhere else in the *N. gonorrhoeae* genome. An in-frame stop codon present in each coding sequence of the sulfite reductase encoding genes, *cysIJ*, within the *N. gonorrhoeae* genome, renders these proteins non-functional [10]. The combination of these deletions and stop codons culminate in a non-functional sulfate reduction pathway in *N. gonorrhoeae.* Unlike *N. gonorrhoeae, S. aureus* is completely lacking any sulfate, thiosulfate or R-sulfonate importers [11, 12]. Not only does it lack the ability to import these important sulfur sources, but similar to *N. gonorrhoeae, S. aureus* lacks the sulfate reductase gene *cysI*, preventing it from reducing sulfate to sulfite [11, 12]. Accordingly, *N. gonorrhoeae* and *S. aureus* share an inability to grow on sulfate as a sole sulfur source, consistent with a non-functional sulfate reduction pathway as described. They are, however, able to use thiosulfate as a sole sulfur source [12, 13], with the L-cysteine requirement of both bacteria being met by thiosulfate, suggesting the ABC transporter complex in *N. gonorrhoeae* is able to import thiosulfate[10], and the presence of a yet unknown thiosulfate uptake system in *S. aureus* [11].

The final step of the pathway, is the biosynthesis of L-cysteine. Cysteine is a key amino acid in proteins, and other biomolecules such as coenzyme A and is required for the synthesis of reducing agents such as thioredoxin and glutathione, which are essential for mitigating intracellular oxidants and resisting host immune responses [8, 10, 14]. One of the primary challenges for bacteria, in particular pathogenic bacteria, is surviving the hostile host environment and evading the host immune system, both of which *N. gonorrhoeae* and *S. aureus* are proficient in [15, 16]. Both pathogens are constantly under pressure of oxidative killing from the host [15, 17, 18], making mechanisms to mitigate this oxidative stress extremely important. The immune system exerts this oxidative killing mechanism primarily through polymorphonuclear neutrophils (PMNs) [15–18]. To mitigate this oxidative stress, an effective intracellular reducing system is required, and most compounds within this system are derived from L-cysteine. Glutathione, synthesised by glutathione reductase, prevents unwanted oxidation of cellular compounds, and regulates the redox balance when encountering free radicals released by the host cell [19, 20]. Thus, intracellular glutathione levels are one of the main oxidative stress mitigating mechanisms deployed during infection. Here is where distinct differences between *N. gonorrhoeae* and *S. aureus* sulfur metabolism can be seen. *N. gonorrhoeae* lacks the ability to recycle glutathione [18] and lacks glutathione import machinery [10], whereas, *S. aureus* only lacks the ability to synthesise glutathione, therefore relying on its import [21].

Bacteria also retain the ability to obtain L-cysteine from the environment. The ability to import L-cysteine and L-cystine is well conserved across many bacterial species, particularly in pathogens, giving a distinct advantage over the energetically expensive *de novo* biosynthesis of L-cysteine [22]. In an oxidative environment such as those of *N. gonorrhoeae* and *S. aureus* infection sites [17, 23], L-cysteine is commonly found in its oxidised form, L-cystine [24]. *N. gonorrhoeae* has ABC transporters for both L-cysteine (*ngo2011-2014*), and L-cystine (*ngo0372-0374*) (Figure 1) [10, 25]. *S. aureus* is also capable of importing L-cysteine and L-cystine using its symporter tcyP (*SA0368*) and ABC transporter tycABC (*SA2202-SA2200*) [12]. However, in the cysteine/cystine deplete or oxidative environment of *N. gonorrhoeae* and *S. aureus* infection, transportation alone would not fulfil L-cysteine requirements, therefore justifying the need for *de novo* biosynthesis of L-cysteine.

L-cysteine biosynthesis occurs in a two-step reaction, catalysed by two enzymes, serine acetyltransferase (SAT) and *O*-acetylserine sulfydrylase (OASS), denoted as CysE and CysK/CysM (OASS isoforms), respectively. Initially CysE, acetylates L-serine, forming *O-*acetylserine and CoA (Figure 2). In the final step, CysK catalyses a condensation reaction of *O-*acetylserine with sulfide, producing L-cysteine (Figure 2).

**Figure 2:**
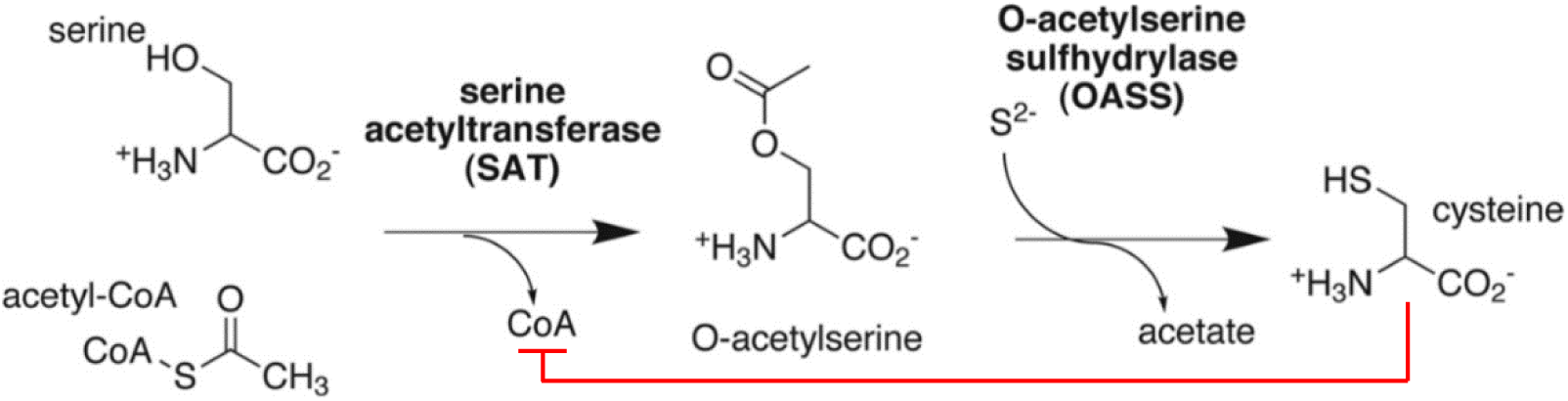
Cysteine biosynthesis pathway. CysE denoted as SAT and CysK denoted as OASS

CysE belongs to the left-handed parallel β-helix family and catalyses the first step of this reaction in both bacteria and plants [27]. In both *S. aureus* and *N. gonorrhoeae,* CysE has been identified as an essential gene [28, 29], and whilst non-essential in *N. meningitidis,* the *cysE* deletion strain confers impaired growth in media [30]. L-cysteine feedback inhibition of CysE is well conserved across bacteria, regulates the first step of the two-step biosynthesis pathway (Figure 2), and acts as the primary method of prevention for high concentrations of intracellular L-cysteine, which are toxic to bacteria [31, 32].

OASS is a PLP dependent enzyme belonging to both the tryptophan synthase β superfamily, and the β-family of pyridoxal 5’-phopshate (PLP) dependent enzymes [5, 32, 33]. There are two commonly found isoforms of OASS, OASS-A (CysK) and OASS-B (CysM) [8]. Most bacteria have both isoforms, using CysK (OASS-A) and CysM (OASS-B) for cysteine biosynthesis using sulfide or thiosulfate respectively. *Salmonella enterica* Servovar typhimurium is one such example, however, expression was found to be environmentally dependent, where CysK (OASS-A) was expressed in excess of CysM (OASS-B) under aerobic conditions, and vice versa under anaerobic conditions [34]. CysM, unlike CysK, can directly use thiosulfate to produce S-sulfocysteine, which is subsequently reduced to L-cysteine [34]. Interestingly, *N. gonorrhoeae* and *S. aureus* have only one OASS isoform, which has higher homology to CysK (OASS-A) and is hypothesised to use sulfide for L-cysteine biosynthesis[10, 35]. With *N. gonorrhoeae* and *S. aureus* lacking the ability to reduce sulfate to sulfide, and relying on thiosulfate for cysteine biosynthesis, a substrate which only the CysM isoform utilises, intriguing questions regarding the role of OASS within these pathogenic bacteria arise, especially considering the annotation of the OASS from *S. aureus* as the CysM isoform [11].

Due to the cytotoxic nature of high L-cysteine concentrations, L-cysteine biosynthesis is well regulated in bacteria. The L-cysteine regulon is controlled at the transcriptional level by CysB, a homotetramer, and a member of the LysR family of transcription factors [10, 36]. CysB can bind to DNA due to the helix-turn-helix motif within the N-terminal domain [37]. The CysE product OAS is unstable and rapidly isomerises to *N-*acetylserine (NAS)[38, 39], both of which bind to CysB, causing positive regulation of L-cysteine biosynthesis and sulfate acquisition genes, however, NAS produces an estimated 15-fold higher response [38, 40, 41]. CysB upregulates the sulfate acquisition genes *cysJIH, cysK,* and *csyP,* whilst negatively autoregulating its own transcription via upstream binding of its own promoter [37, 41, 42]. Pathogenic bacteria depend on their ability to adapt and overcome host defences rapidly, making transcriptional regulation of both the L-cysteine regulon and the sulfate acquisition pathway vital. Interestingly, *cysB* has been identified as an essential gene in *N. gonorrhoeae* [29], but not in *S. aureus* [28].

Feedback inhibition of CysE, and transcriptional regulation are not the only methods of regulating sulfur metabolism and flux. Another mechanism within this pathway lies with the formation of the cysteine synthase complex (CSC). The complex combines the two enzymes that catalyse L-cysteine biosynthesis, CysE and CysK, proven by gel chromatography and fluorescence to consist of one CysE hexamer and two CysK dimers giving a 3:2 protomer ratio [10, 27, 31, 43]. The CSC was first characterised from *S. typhimurium* [43, 44] and there is further evidence of CSC formation in other bacterial species, including *E. coli* [31, 45], *Haemophilus influenzae* [46], and *Mycobacterium tuberculosis* [47]. CSC formation is controlled by the availability of sulfur and involves the C-terminal tail of CysE inserting into the active site of CysK, almost completely inhibiting its activity [48]. When in the complex, CysE loses sensitivity to feedback inhibition by L-cysteine, has reduced substrate inhibition from L-serine and therefore increased catalytic activity [31].

When sulfur is readily available to the cell, the CSC is stabilised by bisulfide, however, with low sulfur availability, OAS accumulates, dissociating the complex and signalling sulfur starvation [31, 49]. This dissociation can occur at µM OAS concentrations (upwards of 50µM OAS) [43, 50], OAS then chemically isomerises to NAS, which binds to transcriptional regulator CysB, promoting expression of sulfate acquisition genes [8, 36, 50]. Fluorescent spectroscopy confirmed binding of the CysE C-terminal peptide binding in the CysK active site, where binding affinity was monitored by CysK activity and PLP fluorescence [4]. The CSC structure has yet to be elucidated, however, there is a crystal structure of CysK from *H. influenzae,* with a CysE 10 residue C-terminal peptide bound in the active site [46]. The key residues for binding were determined via tetrapeptide library screening, indicating a C-terminal isoleucine (Ile) is essential for binding and inhibition of CysK, which remains well conserved across CSC forming bacteria, including *S. typhimurium, H. influenzae* and *E. coli* [31, 46, 51].

## Materials & Methods

### Cloning of Neisseria gonorrhoeae, Staphylococcus aureus, and Escherichia coli cysK for expression in Escherichia coli

The *cysK* genes NGO_0340 (*N. gonorrhoeae*), SAA6008_00518 (*S. aureus*), and NP_416909.1 (*E. coli*) were codon optimised for *E. coli* and ordered from Geneart (ThermoFisher) and Twist Bioscience. The synthetic *cysK* constructs were cloned into expression vector pET28b-PstI with a C-terminal hexahistidine-tag, between PstI and XhoI restriction sites. The plasmids for each *cysK* pET28b plasmid were confirmed by DNA sequencing before transformation into *E. coli* BL21 (DE3) for protein expression. Positive transformants were selected for by growing overnight at 37°C, in Luria-Bertani (LB) agar supplemented with 50 µg.ml^−1^ kanamycin.

### CysK expression and purification

The same method was used for all CysK variants. *E. coli* BL21 (DE3) containing the NGO_0340_pET28b, SAA6008_00518_pET28b, and NP_416909.1_pET28b plasmid were cultured in 1 L LB broth, supplemented with 50 µg.ml^−1^ kanamycin. Cultures were incubated at 37 °C (200 rpm) until an OD_600_ of 0.5-0.7. Protein expression was induced by the addition of 0.75 mM IPTG and cultures were incubated at 22 °C (200 rpm) overnight. Cultures were centrifuged at 4,600 ***g*** for 20 min at 4 °C and the resulting cell pellet was resuspended in lysis buffer (50 mM potassium phosphate pH 7.0, 200 mM NaCl, 20 mM imidazole). One Complete Mini, EDTA-free protease inhibitor tablet (Roche), and 0.1 mM PLP were added prior to cell lysis by sonication. Lysate was centrifuged at 29,097 ***g*** for 25 min at 4 °C and 0.2 µm filtered supernatant was loaded onto a pre-equilibrated HisTrap^TM^ column (GE Healthcare). The column was washed with 20 ml lysis buffer before the elution of CysK using a 50% gradient over 25 ml (50 mM potassium phosphate pH 7.0, 200 mM NaCl, 1 M imidazole).

Immobilised-metal ion affinity chromatography-purified (IMAC-purified) CysK was concentrated at 4 °C using an Amicon^®^ Ultra-15 Centrifugal Filter Units (10 kDa molecular weight cutoff) to a final volume of 5 ml. Concentrated CysK was loaded and run through a HiLoad^TM^ 16/60 Superdex^TM^ 75 column (GE Healthcare), preequilibrated in 50 mM Potassium phosphate pH 7.0, 100 mM NaCl and eluted CysK was collected and stored at 4 °C. Protein concentration was measured by absorbance at 280 nm by Nanodrop^TM^.

### Measuring CysK activity

CysK for kinetic characterisation was purified immediately prior to assays. Enzyme was stored at 4 °C for the duration of the assay, as a rapid decrease in activity was observed when stored at room temperature. Assays were conducted within 14 h post-purification as CysK activity slowly decreased over time. CysK activity was measured by adapting a method from [52]. CysK activity was monitored via absorbance measurements of L-cysteine at 560 nm (*A*_560_) using a SpectraMax ® M Series Multi-Mode Microplate Reader (Molecular devices).

To measure the K_M_ and V_max_ for the substrate *O*-acetylserine (OAS), assays were carried out in 96-well PCR microplates, with a reaction volume of 75 µL, containing variable volumes of 100 mM MOPs, sodium sulfide (Na_2_S) (30 mM for NgCysK without glycerol, 7 mM for NgCysK with glycerol, and 15 mM for SaCysK with and without glycerol), and variable amounts of OAS. The reaction was performed at 37 °C after the addition and subsequent spin down (30 seconds at 2500 ***g***) of 0.2 µg of purified CysK. After addition of acid ninhydrin (100 µl) and glacial acetic acid (100 µl), reaction mixtures were incubated at 95 °C (5 min) forming a pink chromophore (an L-cysteine + ninhydrin derivative), and subsequently incubated on ice (5 min). Ice cold ethanol (100%) was added to 100 µl of each reaction mixture and absorbance of the pink chromophore (560 nm) measured. Enzyme concentration in activity assays was optimised for SaCysK (as SaCysK had noticeably higher reaction rates than NgCysK) by testing various concentrations of SaCysK (0.0058, 0.058, 0.2, 0.4, 0.58, 0.75, 1, 2.5, 3.5, 5.8, and 58 ug; Supplementary Figure 1) and matched by NgCysK. All substrate stocks were prepared in MQ H_2_O. Other components of the reactions and measurement step reagents can be found in the supplementary information. Enzyme working stocks of 0.05 mg.ml^−1^ (1.47 µM SaCysK monomer, 34.011 kDa; 1.47 µM SaCysK monomer, 34.011 kDa, 25% glycerol v/v; 1.48 µM NgCysK monomer, 33.791 kDa; 1.48 µM NgCysK monomer, 33.791 kDa, 25% glycerol v/v) were stored at 4 °C for the duration of assays. The K_M_ and V_max_ were calculated for Na_2_S, by varying the amount of Na_2_S and keeping the concentration of OAS constant at 10 mM across all enzymes. K_M_ and V_max_ values were determined by non-linear regression fit of the Michaelis Menten (Equation 1), allosteric sigmoidal (Equation 2), substrate inhibition (Equation 3), or the allosteric sigmoidal with substrate inhibition equation (Equation 4) using GraphPad Prism (GraphPad Software Version 10.2.3).

The initial velocity of the reaction was derived from linear-regression analysis of the first 5 minutes of the reaction using a cysteine standard curve (Equation 1). All concentrations were collected in triplicate. Cysteine produced was calculated using a cysteine standard curve equation (Equation 5), and k_cat_ was calculated by dividing the V_max_ rate (M.s^−1^) by the molar concentration of dimeric enzyme. The catalytic efficiency k_cat_/K_M_ (M^−1^s^−1^) was calculated by dividing the k_cat_ (s^−1^) by the K_M_ (M). Catalytic efficiency values were calculated for each substrate.

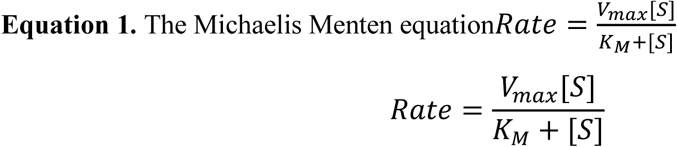

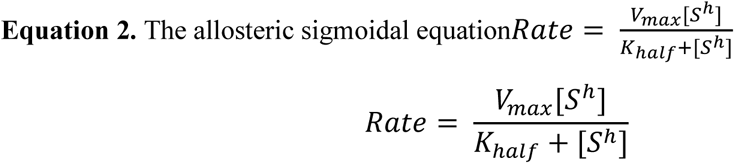

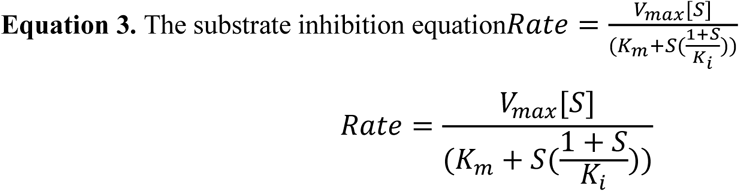

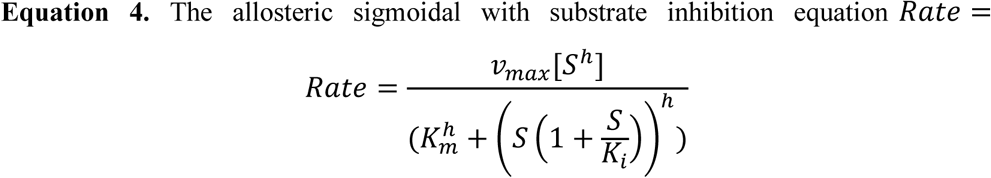

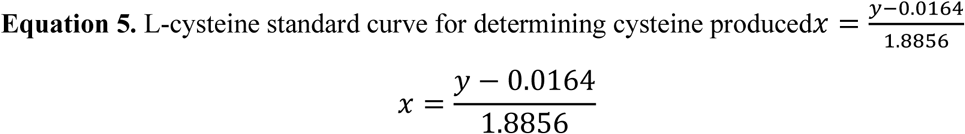

Thiosulfate substrate assays were conducted using a modified method from above. The key modification was substitution of thiosulfate for sodium sulfide in the reaction. Thiosulfate was added at low to high varied concentrations (final reaction concentration, 1-50 mM).

### Small angle X-ray scattering (SAXS) analysis of CysK and formation of the cysteine synthase complex

#### Purification of CysE

NgCysE and EcCysE were purified using IMAC and gel filtration chromatography with minor modifications. CysE pellets were resuspended in lysis buffer (50 mM Tris pH 8.0, 200 mM NaCl, 20 mM imidazole). No PLP was added prior to sonication. Lysate was centrifuged at 20000 ***g*** for 20 min at room temperature. The pre-equilibrated HisTrap^TM^ column (GE Healthcare) was washed with 20 ml lysis buffer before the elution of CysE using a 50% gradient over 25 ml (50 mM Tris pH 8.0, 200 mM NaCl, 1 M imidazole) [53]. IMAC-purified CysE was concentrated at 15 °C using an Amicon^®^ Ultra-15 Centrifugal Filter Units (10 kDa molecular weight cutoff) to a final volume of 0.75 ml [53]. Concentrated CysE was run through an Enrich 650 analytical gel filtration column (BioRad), pre-equilibrated in 50 mM Tris pH 8.0, 100 mM NaCl and stored at room temperature.

#### Preparation of samples

NgCysK, EcCysK, SaCysK, NgCysE, and EcCysE were expressed and purified as mentioned above. SaCysE was unable to be purified as it remained insoluble. All CysK samples were concentrated using Amicon^®^ Ultra-15 Centrifugal Filter Units (10 kDa molecular weight cutoff, centrifuged at 4 ⁰C and 3000 ***g***) and made up with glycerol (25% v/v final) and transported on ice. All CysE samples were prepared as close to the time of departure as possible (one or two days beforehand) and transported at room temperature to avoid cold inactivation [54] or aggregation due to overconcentration. CysE samples were concentrated on site at the Australian Synchrotron using Amicon^®^ Ultra-0.5 Centrifugal Filter Units (10 kDa molecular weight cutoff, centrifuged at 4 ⁰C and 13,000 ***g***).

#### Data Collection

Experiments were conducted using the Australian Synchrotron SAXS/WAXS beamline equipped with a Dectris PILATUS 1M/200k detector coupled to a size exclusion chromatography (SEC) with a sheath flow cell [55]. Protein samples not analysed for CSC formation (non-CSC formation) were prepared in Potassium phosphate buffer (50 mM Potassium phosphate pH 7.0, 100 mM NaCl), and protein samples for potential CSC formation analysis were prepared in TRIS buffer (50 mM Tris pH 8.0, 100 mM NaCl). Non-CSC formation samples were prepared to final concentrations of 4, 6 and 8 mg/ml in a 96-well plate (45 µl total volume). This allowed for correction of concentration dependent behaviours and the mitigation of potential sample aggregation. Potential CSC forming samples were prepared to a 3:2 CysE:CysK monomeric ratio with a 6 mg/ml final concentration in a 96-well plate (60 µl total volume). SEC was conducted using either a 3.2 mL Superdex S200 5/150 (Cytiva) for individual CysK non-CSC formation samples, or a 24 mL Superdex S200 5/150 (Cytiva) for CysK and CysE samples analysed for CSC formation, respectively. Each column was equilibrated with the appropriate buffer prior to sample loading; Potassium phosphate for the 3.2 mL Superdex S200 5/150 (Cytiva) column (for non-CSC samples); and TRIS (pH 8.0) for the 24 mL Superdex S200 5/150 (Cytiva) column (for potential CSC forming samples).

Scattering images were collected using instrument parameters as seen in Table 1 [56]. Scattering intensity was measured as a function of momentum transfer (*q*), where *q* = 4πsin(θ)/λ and 2θ is the scattering angle in Å [57]. Samples (45 or 60 µl injection volume) were run through the size exclusion columns at a flow rate of 0.4 ml/min (Table 2 and into a 1.5 mm thin-walled glass capillary where they were exposed to continuous one second x-ray bursts over an ∼10 (3.2 ml column) or ∼60 (24 ml column) minute elution period.

**Table 1.** SEC-SAXS Data Collection Parameters.

|  |  |
| --- | --- |
| **Instrument** | Australian Synchrotron SAXS/WAXS beamline with Dectris PILATUS 1M/200k detector [56] |
| **Wavelength ($\text{\AA}$)** | 1.033 |
| **Beam size ($\mu\text{m}$)** | 250 x 450 |
| **Camera length (m)** | 2.683 |
| **$q$ measurement range ($\text{\AA}^{-1}$)** | 0.0015-3.0 |
| **Absolute scaling method** | Comparison with scattering from 1 mm pure H<sub>2</sub>O |
| **Normalisation** | To transmitted intensity by beam-stop counter |
| **Monitoring for radiation damage** | X-ray dose maintained below 210 Gy, data frame-by-frame comparison |
| **Exposure time** | Continuous 1s data-frame measurements of SEC elution |
| **Sample configuration** | SEC-SAXS with sheath-flow cell [55], effective sample path length 0.49 mm |
| **Sample temperature (°C)** | 8 |

**Table 2.** SAXS sample details.

|  | NgCysK | EcCysK | SaCysK | NgCysE | EcCysE | NgCSC | EcCSC |
| --- | --- | --- | --- | --- | --- | --- | --- |
| **Organism** | *Neisseria gonorrhoeae* | *Escherichia coli* | *Staphylococcus aureus* | *Neisseria gonorrhoeae* | *Escherichia coli* | *Neisseria gonorrhoeae* | *Escherichia coli* |
| **Source** | *E. coli* expressed | *E. coli* expressed | *E. coli* expressed | *E. coli* expressed | *E. coli* expressed | *E. coli* expressed | *E. coli* expressed |
| **UniProt sequence ID (residues in construct)** | Q5F9Q2 (1-310) + C-terminal Hexa His-tag | P0ABK5 (1-323) + C-terminal Hexa His-tag | A0A0U1ME23 (1-301) + C-terminal Hexa His-tag | A0A1D3FX11 (1-272) + N-terminal Hexa His-tag | P0A9D4 (1-273) + N-terminal Hexa His-tag | Q5F9Q2 (1-310) C-terminal Hexa His-tag + A0A1D3F (1-272) N-terminal Hexa His-tag | P0ABK5 (1-323) C-terminal Hexa His-tag + P0A9D4 (1-273) N-terminal Hexa His-tag |
| **Extinction coefficient [$A_{280}$, 0.1% (w/v)]** | 0.594 | 0.535 | 1.112 | 0.598 | 0.855 | - | - |
| **$\bar{v}$ from chemical composition (cm<sup>3</sup> g<sup>-1</sup>)** | 0.743 | 0.743 | 0.733 | 0.740 | 0.739 | - | 0.741 |
| **Particle contrast from sequence and solvent constituent, $\Delta\rho$ ($\rho_{\text{protein}} - \rho_{\text{solvent}}$; 10<sup>10</sup> cm<sup>-2</sup>)** | 2.775 | 2.780 | 2.895 | 2.799 | 2.811 | - | 2.798 |
| ***M* from chemical composition (Da)** | 32726 | 36780 | 32946 | 31600 | 31600 | 320504 | 336720 |
| **SEC-SAXS column (3.2 mL Superdex S200 5/150 (Cytiva))** |  |  |  |  |  |  |  |
| **Loading concentration (mg.ml<sup>-1</sup>)** | 6 | 6 | 6 | 6 | - | - | - |
| **Injection volume (ul)** | 45 | 45 | 45 | 45 | - | - | - |
| **Flow rate (ml.min<sup>-1</sup>)** | 0.4 | 0.4 | 0.4 | 0.4 | - | - | - |
| **SEC-SAXS column (24 mL Superdex S200 5/150 (Cytiva))** |  |  |  |  |  |  |  |
| **Loading concentration (mg.ml<sup>-1</sup>)** | - | 6 | - | - | 6 | 6 | 6 |
| **Injection volume (ul)** | - | 45 | - | - | 45 | 60 | 60 |
| **Flow rate (ml.min<sup>-1</sup>)** | - | 0.4 | - | - | 0.4 | 0.4 | 0.4 |
| **Solvent** | 50 mM TRIS, 100 mM NaCl, 0.1% Azide, pH 8.0 |  |  |  |  |  |  |

#### Data Processing

The beamline intensity normalisation and subsequent data reduction was performed using the Australian Synchrotron developed software scatterBrain (version 2.71) (Table 3). Scattering files produced were aligned with the inline SEC profile using CHROMIXS [58]. The sample-scattering frames were selected from the largest peak, corresponding with the dimeric or hexameric protein for all CysK and CysE enzymes respectively. In CSC samples, sample-scattering frames were selected from the initial peak corresponding to the decameric complex. In all cases, the buffer-scattering region was selected from upstream of the first protein elution peak. The buffer-scattering was subtracted from the selected sample-scattering and the processed scattering profiles were exported as *.dat files which are readable by the ATSAS package programmes. Unless stated otherwise, all downstream data processing was performed using the ATSAS program suite (version 3.1.0) [59].

**Table 3.**
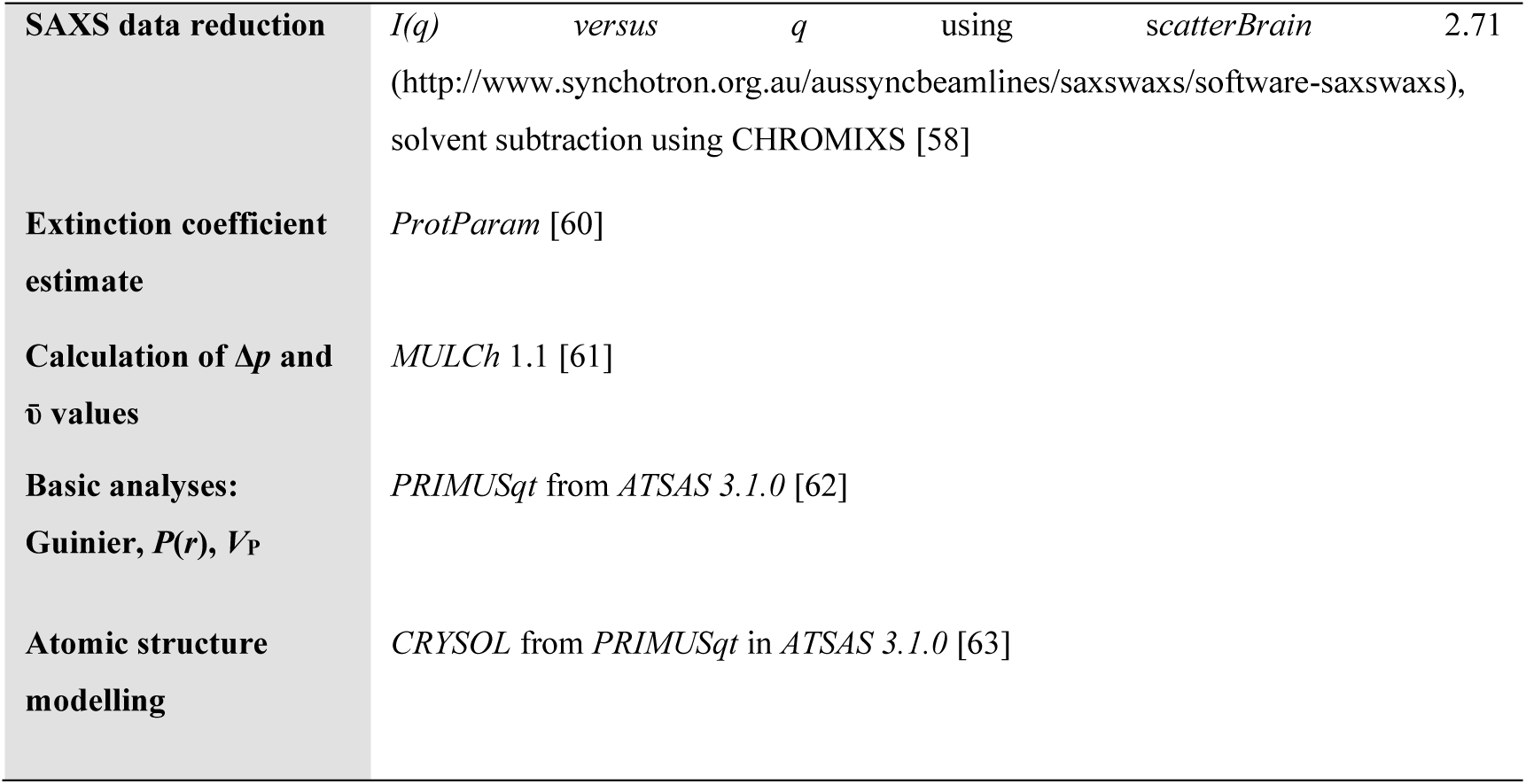
Reduction and analysis software.

Subtracted scattering data was analysed using the PRIMUSqt software [59, 62]. Within PRIMUSqt, the “Radius of Gyration” and the “Distance Distribution” analysis functions were both used to determine the radius of gyration (R_g_) and the relative intensity (forward scattering intensity, I(0)) for each sample. The “Radius of Gyration” function uses the Guinier Approximation of low-resolution scattering data (the low-*q* region) to calculate reciprocal space values. Keeping in line with the standard criteria for globular proteins, the plotted range of Guinier plots was kept below *q*R_g_ = 1.3 [57, 64–66]. These plots were assessed for quality of fit using the fidelity of the model (ranging from 0.0 being the lowest in quality, to 1.0 being the highest quality) alongside the random distribution of residuals averaging around zero. Aggregation of the protein samples was evaluated based on the linearity of the Guinier plots (non-linear regions indicate aggregation). The “Distance Distribution” uses the pair distance distribution function (P(r)) to calculate the maximum particle dimensions (D_max_), the real-space R_g_, and I(0). Interatomic distance distributions (r) are calculated using GNOM [67] from the “Distance Distribution” function. D_max_ was defined by the value of r where the P(r) curve reaches zero. This distance distribution data was fit to an approximate maximum *q* of 0.3 Å^−1^ (Tables 1, 2, 3, and Supplementary Figure 3). D_max_ was manually adjusted to the lowest possible value whilst still producing a smooth curve, with P(r)>0 and a random distribution of residuals around zero. The P(r) curves were normalised by dividing by I(0) (from the “Guinier analysis”) to remove any confounding effects of concentration.

The “Porod Volume” function in PRIMUSqt was used to calculate the Porod Volume (V_P_), using a *q*-range limited to the Porod-Debye region and reciprocal space values determined by the “Radius of Gyration” function. The Porod-Debye region plotted was defined by the scattering data that reached and maintained a plateau within the Porod plot (Supplementary Figure 3) [68]. This subset of data was used to determine the V_p_ which in turn was used together with calculated molecular weights (MW) to determine the particle densities (d_particle_) for each CysK variant in each condition tested. The MW was determined using the protein sequence analysis programme ProtParam [60]. The d_particle_ was calculated using Equation 6, where MW is the molecular weight of the protein and V_p_ is the calculated porod volume [68].

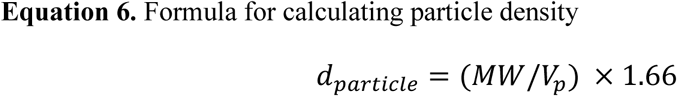

Dimensionless Kratky plots were created by multiplying *q* by the R_g_ (from the “Guinier analysis”) to remove effects of protein size and molecular weight. The intensity was normalised by dividing by I(0) (from the “Guinier analysis”) to remove any confounding effects of concentration.

Theoretical scattering of known enzyme structures overlayed onto experimental scattering data plots were created using “CRYSOL online” from ATSAS online 3.2.1 web services tool. All CysK enzymes experimental scattering data were overlaid with the theoretical scattering of our *N. gonorrhoeae* CysK crystal structure (NgCysK, 9NLD) from the PDB. All CysE enzymes experimental scattering data were overlaid with the theoretical data of their corresponding organisms’ crystal structure from the PDB (NgCysE with 6WYE, and EcCysE with 1T3D). Crysol was used to validate the enzymes and determine similarities between enzymes in solution (SAXS) and rigid models (X-ray crystallography).

### Cysteine synthase complex formation

The protocol for forming the cysteine synthase complex (CSC) was based on a method by [31].

#### Monitoring cysteine synthase complex formation by gel filtration chromatography

A 1 ml solution of NgCysE 6 µM: NgCysK 4 µM (monomer ratios) was manually injected into the injection loop of the FPLC for injection into the Enrich 650 analytical gel filtration column (Bio-Rad Laboratories, USA), preequilibrated in gel filtration buffer (50 mM Potassium phopshate pH 7.0, 100 mM NaCl). After sample injection, the column was washed with 28 ml of gel filtration buffer (50 mM Potassium phosphate pH 7.0, 100 mM NaCl). Fractions were collected and run on a 12% SDS PAGE gel. This was repeated with a NgCysE 4 µM: NgCysK 0.05 µM 1 ml solution, without fractions being run on a 12% SDS PAGE gel. The same method was repeated using a EcCysE 6 µM: EcCysK 4 µM monomeric ratio.

#### Monitoring cysteine synthase complex formation by small angle X-ray scattering

NgCysK, EcCysK, NgCysE, and EcCysE were expressed and purified and prepared as mentioned above. Data was processed in the same manner as discussed above with particular emphasis on the scattering plots, which display the key differences between CysK, CysE and CSC scattering.

## Results and Discussion

### Substrate specificity of CysK from two bacterial pathogens lacking the sulfate reduction pathway

Given the importance of sulfur metabolism, and its central role in L-cysteine biosynthesis in bacteria, investigating the kinetic function of the final step of L-cysteine biosynthesis in these two pathogenic bacteria lacking the sulfate reduction pathway is vital. Kinetic analyses of CysK from *N. gonorrhoae* and *S. aureus* were conducted as we endeavour to elucidate the differences in substrate utilisation and in turn sulfur metabolism (Figure 1). The production of L-cysteine by CysK was determined by measuring the absorbance of a pink chromophore that forms when ninhydrin interacts with L-cysteine [69]. It is well documented that addition of 25% (v/v) glycerol (final concentration) significantly improves stability of the enzyme, mitigating decreases in activity caused by freeze/thawing [70]. Glycerol has been used on many occasions for CysK stabilisation during storage [71, 72], resuspension of pellets and samples [73, 74], elution, and dialysis [31, 73]. However, we discovered the presence of glycerol had a significant effect on the kinetics of NgCysK and SaCysK (Supplementary Figure 4, Supplementary Figure 5, Supplementary Table 1, and Supplementary Table 2). Therefore, kinetic analyses presented here were completed in the absence of glycerol, using freshly purified enzyme.

The kinetic parameters of NgCysK for substrate OAS (Table 3) were calculated from fitting an Allosteric sigmoidal model (R^2^ = 0.9702) of rate versus OAS (Figure 3A). The overall fit for the Allosteric sigmoidal equation is good, giving a K_half (OAS)_ of 1.541 mM, a k_cat (OAS)_ of 1.166 x 10^6^ s^−1^, and a k_cat_/K_M (OAS)_ of 7.570 x 10^8^ M.s^−1^ for the dimer (Table 4). A saturating concentration of 30 mM Na_2_S was used for the collection of the Allosteric sigmoidal plot. The kinetic parameters of NgCysK for substrate Na_2_S were calculated from fitting an Allosteric sigmoidal model (R^2^ = 0.9088) of rate versus Na_2_S (Figure 3B). The overall fit for the Allosteric sigmoidal equation is good, giving a K_half (Na2S)_ of 23.620 mM, a k_cat (Na2S)_ of 1.345 x 10^6^ s^−1^, and a k_cat_/K_M (Na2S)_ of 5.700 x 10^7^ M.s^−1^ for the dimer (Table 4).

**Figure 3:**
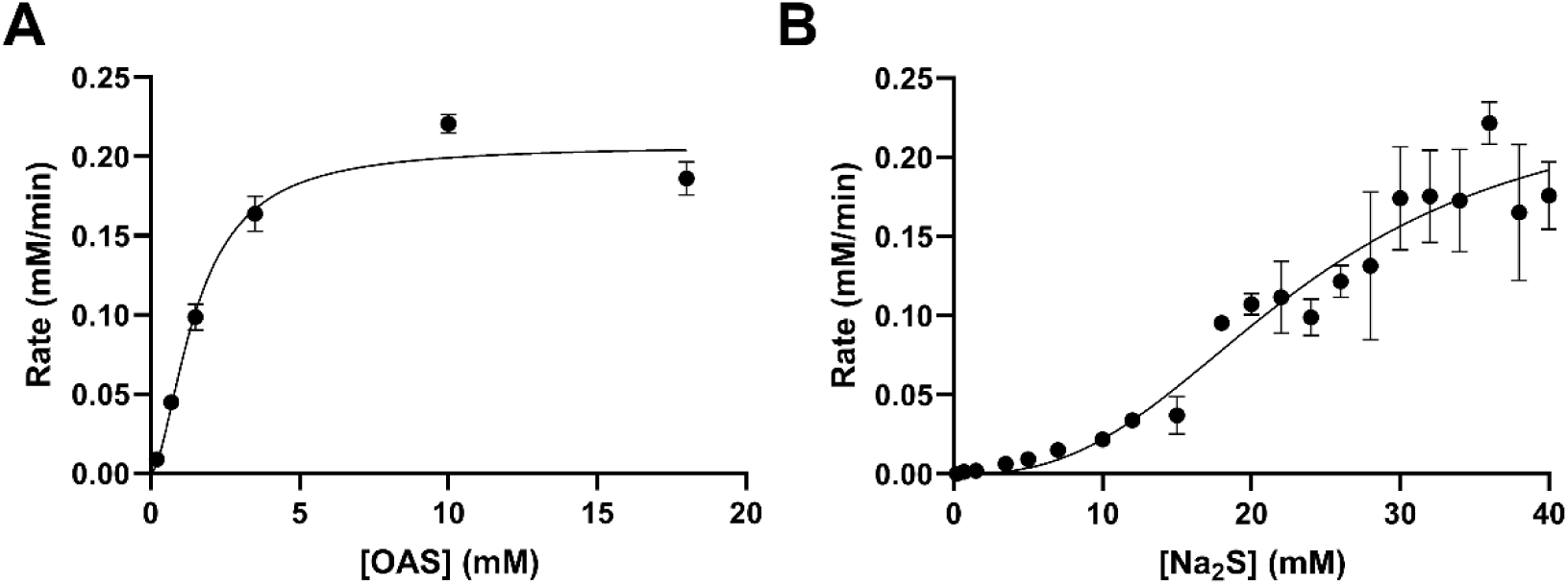
Kinetic analysis of NgCysK substrates OAS and Na_2_S in the absence of glycerol. (A) Allosteric sigmoidal fit (occuring when the enzyme has cooperative subunits) for OAS. (B) Allosteric sigmoidal fit for Na_2_S. OAS and Na_2_S assays were collected at saturating concentrations of 10 mM OAS and 30 mM Na_2_S, respectively. Plotted data points represent mean alongside SEM of three replicates.

**Table 4:** Kinetic parameters of NgCysK in the absence of glycerol.

| Parameter | *O*-acetylserine (OAS)<sup>a</sup> | Sodium sulfide (Na<sub>2</sub>S)<sup>a</sup> |
| --- | --- | --- |
| $v_{\max}$ (mM.min<sup>-1</sup>) | 0.207 ± 0.016 | 0.239 ± 0.111 |
| $K_{\text{half}}$ (mM) | 1.541 ± 0.306 | 23.620 ± 10.780 |
| $k_{\text{cat}}$ (s<sup>-1</sup>)<sup>b</sup> | (1.166 ± 0.091) × 10<sup>6</sup> | (1.345 ± 0.625) × 10<sup>6</sup> |
| $k_{\text{cat}}/K_M$ (M.s<sup>-1</sup>) | (7.570 ± 1.616) × 10<sup>8</sup> | (5.700 ± 3.710) × 10<sup>7</sup> |
| $h$ | 1.725 ± 0.513 | 2.669 ± 1.086 |
| $R^2$ | 0.9702 | 0.9088 |

<sup>a</sup> Error is SEM of three replicates
<sup>b</sup> $k_{\text{cat}}$ calculated by dividing the rate (M.s<sup>-1</sup>) by enzyme concentration using the concentration of the NgCysK dimer (33.791 kDa)

The kinetic parameters of SaCysK for substrate OAS (Table 5) were calculated from fitting a Substrate inhibition model (R^2^ = 0.9301) of rate versus OAS (Figure 3A). The overall fit for the Substrate inhibition equation is good, giving a K_M (OAS)_ of 1.378 mM, a k_cat (OAS)_ of 7.155 x 10^5^ s^−1^, and a k_cat_/K_M (OAS)_ of 5.194 x 10^8^ M.s^−1^ for the dimer (Table 5). A saturating concentration of 15 mM Na_2_S was used for the collection of the Substrate inhibition plot. The kinetic parameters of SaCysK for substrate Na_2_S were also calculated from fitting a Substrate inhibition model (R^2^ = 0.6206) of rate versus Na_2_S (Figure 3B). The overall fit for the Substrate inhibition equation is poorer than that of OAS due to the error at 12 and 15 mM Na_2_S concentrations, alongside the low rate at 8 mM Na_2_S. However, kinetic parameters were obtained from the data, giving a K_M (Na2S)_ of 2.306 mM, a k_cat (Na2S)_ of 3.081 x 10^5^ s^−1^, and a k_cat_/K_M (Na2S)_ of 1.336 x 10^8^ M.s^−1^ for the dimer (Table 5).

**Table 5:** Kinetic parameters of SaCysK in the absence of glycerol.

| Parameter | *O*-acetylserine (OAS)<sup>a</sup> | Sodium sulfide (Na<sub>2</sub>S)<sup>a</sup> |
| --- | --- | --- |
| $v_{max}$ (mM.min<sup>-1</sup>) | $0.126 \pm 0.032$ | $0.054 \pm 0.034$ |
| $K_M$ (mM) | $1.378 \pm 0.592$ | $2.306 \pm 3.573$ |
| $k_{cat}$ (s<sup>-1</sup>)<sup>b</sup> | $(7.155 \pm 1.828) \times 10^5$ | $(3.081 \pm 1.950) \times 10^5$ |
| $k_{cat}/K_M$ (M.s<sup>-1</sup>) | $(5.194 \pm 2.594) \times 10^8$ | $(1.336 \pm 2.236) \times 10^8$ |
| $K_i$ (mM) | $6.265 \pm 2.489$ | $15.970 \pm 12.793$ |
| $R^2$ | 0.9301 | 0.6206 |

<sup>a</sup> Error is SEM of three replicates
<sup>b</sup> $k_{cat}$ calculated by dividing the rate (M.s<sup>-1</sup>) by enzyme concentration using the concentration of the SaCysK dimer (34.011 kDa)

**Figure 3:**
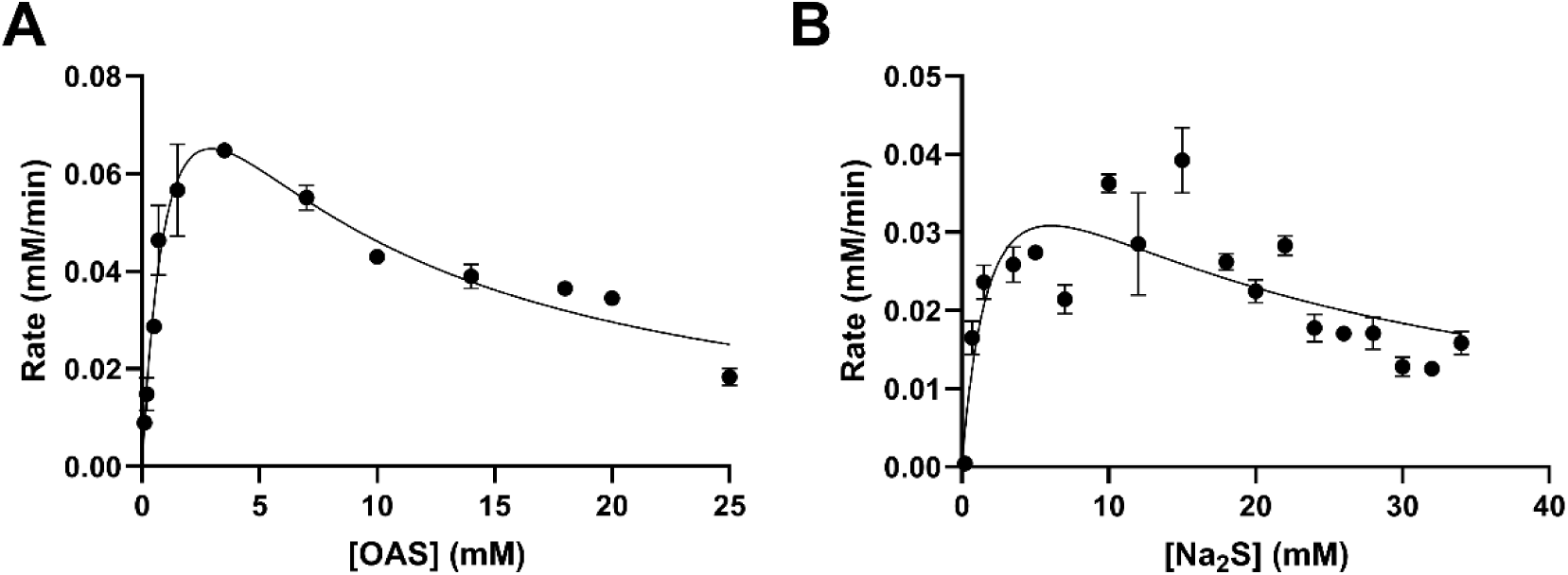
Kinetic analysis of SaCysK substrates OAS and Na_2_S in the absence of glycerol. (A) Substrate inhibition fit (occurs when the enzyme exhibits inhibition at high substrate concentrations) for OAS. (B) Substrate inhibition fit for Na_2_S. OAS and Na_2_S assays were collected at saturating concentrations of 10 mM OAS and 15 mM Na_2_S, respectively. Plotted data points represent mean alongside SEM of three replicates.

The difference in K_half_ and K_M_ values for each substrate in NgCysK (1.541 mM OAS versus 23.620 mM Na_2_S) and SaCysK (1.378 mM OAS versus 2.306 mM Na_2_S), respectively, indicates a greater affinity for OAS compared with Na_2_S in both enzymes. Interestingly, this affinity is similar to the plant *O-*acetylserine sulfhydrylase isozyme A and C from *Datura innoxia* [72], but not to other bacterial homologues [34, 71, 75]. However, the earlier studies were conducted in the presence of glycerol, which we have determined to largely affect the kinetics of each substrate. The specificity constants, k_cat_/K_M_, are 7.57 x 10^8^ (OAS) and 5.700 x 10^7^ M.s^−1^ (Na_2_S) for NgCysK, and 5.194 x 10^8^ (OAS) and 1.336 x 10^8^ M.s^−1^ (Na_2_S) for SaCysK. Catalytic efficiency (k_cat_/K_M_) values of ≥10^8^ M.s^−1^ indicate the enzyme’s reaction rate is diffusion limited [76], and in the case of NgCysK the reaction rate is limited by diffusion of OAS (k_cat_/K_M:_ 7.57 x 10^8^ M.s^−1^), but not Na_2_S (k_cat_/K_M:_ 5.700 x 10^7^ M.s^−1^). For SaCysK the reaction rate is limited by diffusion of both substrates, OAS (k_cat_/K_M:_ 5.194 x 10^8^ M.s^−1^), and Na_2_S (k_cat_/K_M:_ 1.336 x 10^8^ M.s^−1^).

There is a ∼13.3-fold increase in k_cat_/K_M_ for OAS compared to Na_2_S attributable to NgCysK having a ∼15.33-fold increased affinity for OAS (K_half_: 1.541 mM) compared to Na_2_S (K_half_: 23.620 mM). In SaCysK, there is a ∼3.89-fold increase in k_cat_/K_M_ for OAS compared to Na_2_S attributable to SaCysK having a ∼1.67-fold increased affinity for OAS (K_M_: 1.378 mM) compared to Na_2_S (K_M_: 2.306 mM). Intriguingly, the kinetics parameters for substrates OAS and Na_2_S vary extensively across CysK homologues. For OAS, positive cooperativity is seen at low concentrations with substrate inhibition above 25 mM in the plant species *D. innoxia* [72]. Plant species *Phaseolus vulgaris* and *Phaseolus polyanthus* exhibit Michaelis-Menten kinetics [71], and *S. typhimurium* exhibits substrate inhibition above 7.5 mM [34, 75]. For NgCysK there is an allosteric sigmoidal relationship to OAS with positive cooperativity at low concentrations and an eventual plateau beginning at ∼5 mM. In comparison SaCysK displays a strictly substrate inhibition relationship, beginning to drop after ∼5 mM OAS. For Na_2_S, substrate inhibition in *P. vulgaris* is observed after reaching a maximal rate at ∼1.5 mM S^2−^ with positive cooperativity (an allosteric sigmoidal model) observed at concentrations below 0.3 mM [71]. Similarly, *D. innoxia* OASS (isozyme C) exhibits positive cooperativity below ∼0.18 mM S^2−^ with substantial substrate inhibition after 0.2 mM [72]. *P. polyanthus* CysK shows no allosteric sigmoidal (positive cooperativity) response to S^2−^, however, fits a substrate inhibition model [71], which is also found in *S. typhimurium* after 0.25 mM S^2−^ [75], similar to the response seen in SaCysK after ∼8 mM Na_2_S. For NgCysK, only an allosteric sigmoidal response is seen with positive cooperativity at low concentrations and a slight plateau beginning at ∼30 mM. It is intriguing to see such variable relationships of the CysK enzyme with its substrates, and of particular interest is the stark difference we see between SaCysK and NgCysK, two enzymes from organisms lacking the same sulfate reduction pathway (Figure 1). Based on the reaction mechanism of CysK homologues [77, 78], and their shared substrates, we predict that both NgCysK and SaCysK function through a bi-bi ping-pong mechanism, in keeping with the proposed alternating cycle of binding and releasing each substrate, being unable to have both substrates bound at the same time.

Both *S. aureus* and *N. gonorrhoeae* lack the sulfate reduction pathway. However, with their ability to grow on media with thiosulfate as the only sulfur source, this raised questions regarding the functionality and characterisation of the CysK enzyme in both *S. aureus* and *N. gonorrhoeae*. The CysM isoform can use both sulfide and thiosulfate as sulfur donors during L-cysteine biosynthesis, therefore, although the kinetic parameters fit with other CysK enzymes, we checked for dual substrate usage, especially due to the previous annotation of the OASS from *S. aureus* as the CysM isoform [11]. To establish both enzymes as a CysK isoform of OASS, we conducted the same kinetic assays as above, replacing Na_2_S with thiosulfate (S_2_O_3_^2−^). Non-linear regression fit of a Michaelis-Menten model gave a very poor fit to the data (R^2^ = 0.2058 (NgCysK) and 0.1426 (SaCysK)) resulting in a v_max_ and K_M_ of 0.002 ± 0.003 mM.min^−1^ and 5.780 ± 11.476 mM, respectively, for NgCysK (Figure 4, Table 6), and 0.000 ± 373,758.5 mM.min^−1^ and 7.727 ± ∞ mM, respectively, for SaCysK (Table 6). The error present in the K_M_ of both enzymes is greater than the K_M_, taking the K_M_ below 0 mM (Table 6). This, alongside the exceptionally low V_max_, which lays outside the values measured, indicates there is no enzyme activity detected for any thiosulfate concentration, therefore, establishing these enzymes from both *S. aureus* and *N. gonorrhoeae* are in fact CysK isoforms, incapable of utilising thiosulfate as a sulfur donor for the synthesis of cysteine.

**Figure 4:**
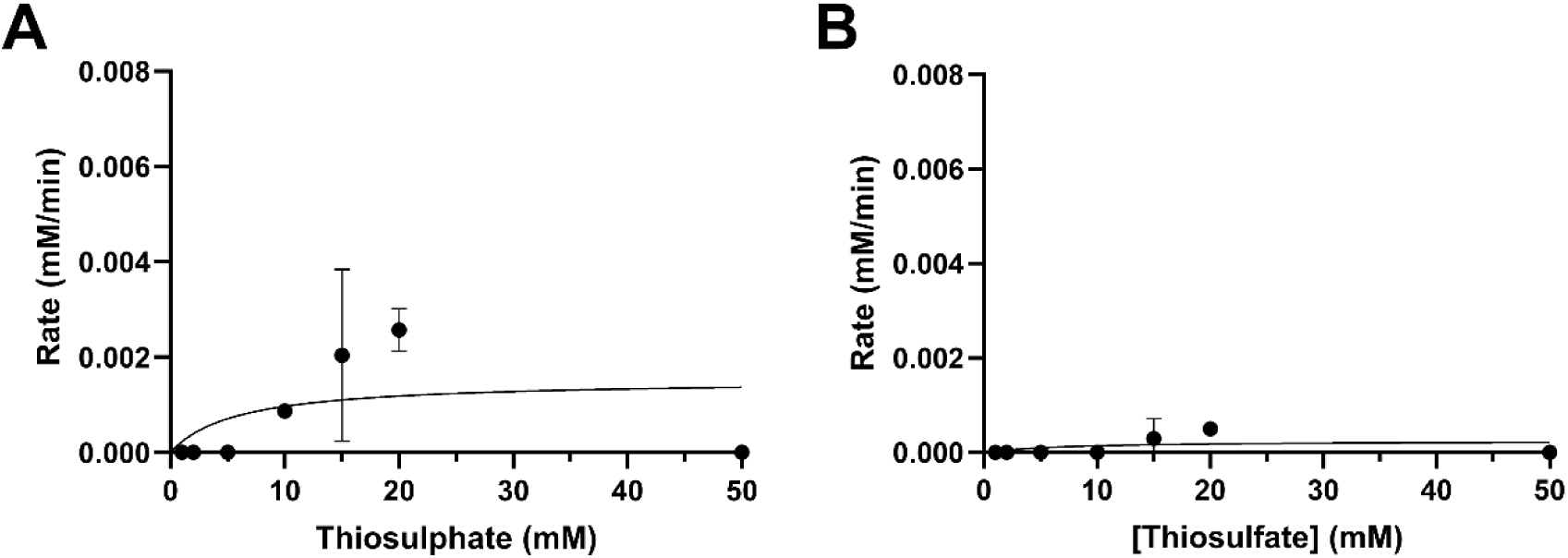
Thiosulfate activity assays for NgCysK and SaCysK. Michaelis-Menten model is fit to both data sets. (A) NgCysK thiosulfate activity. (B) SaCysK thiosulfate activity. Both assays were conducted at saturating concentrations of OAS (10 mM).

**Table 6:** Kinetic parameters of thiosulfate for NgCysK and SaCysK.

| Parameter | NgCysK<sup>a</sup> | SaCysK<sup>a</sup> |
| --- | --- | --- |
| $V_{\max}$ (mM.min<sup>-1</sup>) | 0.002 ± 0.003 | 0.000 ± 373,758.5 |
| $K_M$ (mM) | 5.780 ± 11.476 | 7.727 ± ∞ |
| $k_{\text{cat}}$ (s<sup>-1</sup>)<sup>b</sup> | (8.589 ± 16.300) × 10<sup>3</sup> | (1.417 ± 2.117 × 10<sup>12</sup>) × 10<sup>3</sup> |
| $k_{\text{cat}}/K_M$ (M<sup>-1</sup>.s<sup>-1</sup>) | (1.486 ± 4.080) × 10<sup>6</sup> | (1.834 ± ∞) × 10<sup>5</sup> |
| $R^2$ | 0.2058 | 0.1426 |

<sup>a</sup> Error is SEM of two replicates
<sup>b</sup> $k_{\text{cat}}$ calculated by dividing the rate (M.s<sup>-1</sup>) by enzyme concentration using the concentration of the CysK dimer (NgCysK: 33.791 kDa; SaCysK: 34.011 kDa)
∞ Infinity symbol indicates error was so high it could not be calculated and is therefore infinity

### Small angle X-ray scattering shows no conformational change in the presence of O-acetylserine

Size exclusion chromatography (SEC) combined with small angle X-ray scattering (SEC-SAXS) analysis [55] was performed on CysK enzymes from both, *N. gonorrhoeae* and *S. aureus,* two organisms lacking the sulfate reduction pathway, to determine the effect of OAS binding to both NgCysK and SaCysK in solution. Throughout our analysis, we used the well-characterised CysK enzyme from *E. coli* as a comparison to a “true” CysK [7, 44, 54, 79–81]. The scattering data of NgCysK and SaCysK with and without OAS, and the EcCysK, indicated a folded globular protein (Figure 5), and the calculated molecular mass (Qp, MoW Fischer method, Vc), combined with the pair wise distribution (P(r)) analysis (Figure 5G) support a dimer of CysK monomers for all organisms. However, when comparing MW calculations of all four methods, we see the Porod volume MW calculation consistently gives a 1.39-1.66-fold higher molecular weight estimation for all CysK enzymes, including both NgCysK and SaCysK, in the presence of OAS (Tables 7 and 8). This overestimation can occur due to two limitations of this particular method, a large hydration shell as a result of proteins surface residues interacting with water molecules in solution, and high flexibility of particular regions [68, 82]. The C-terminal region of the CysK enzymes alongside their active site regions, are particularly flexible, combined with the hydration shell of the enzymes in solution may have led to the overestimation of the molecular weight.

**Figure 5:**
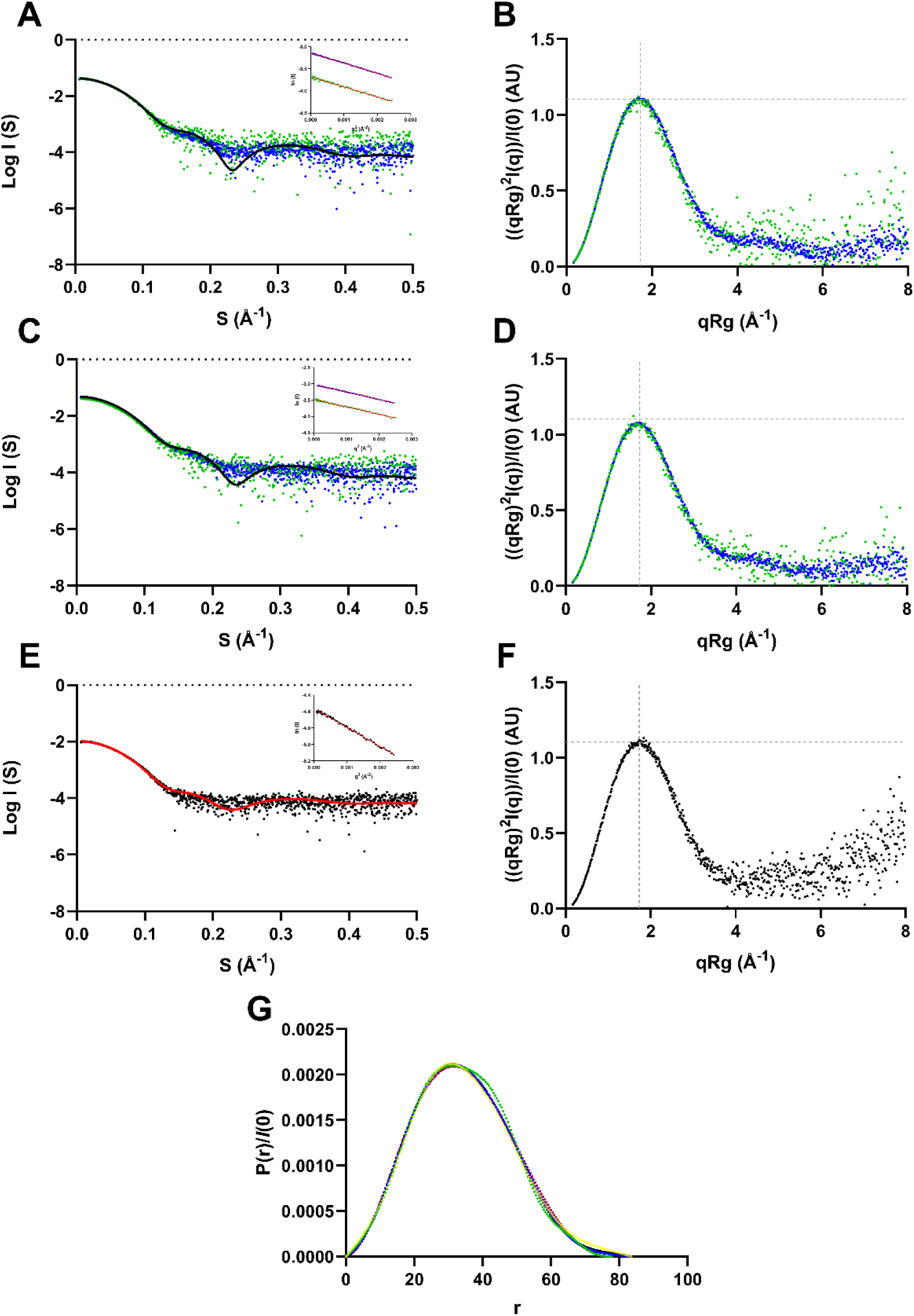
SAXS profiles of NgCysK and SaCysK in the presence and absence of OAS, and EcCysK. (A) Scattering profiles of NgCysK in the absence (blue circles) and presence (green circles) of 10 mM OAS. (B) Kratky plot of NgCysK in the absence (blue circles) and presence (green circles) of 10 mM OAS. (C) Scattering profiles of SaCysK in the absence (blue circles) and presence (green circles) of 10 mM OAS. (D) Kratky plot of SaCysK in the absence (blue circles) and presence (green circles) of 10 mM OAS. (E) Scattering profile of EcCysK. Experimental EcCysK scattering shown in black. Theoretical scattering (red line) calculated from NgCysK crystal structure (9NLD). (F) Kratky plot of EcCysK. (G) Pair distance distribution function (P(r)) analysis of NgCysK and SaCysk in the absence and presence of OAS, and EcCysK. NgCysK shown in black; NgCysK + OAS shown in green; SaCysK shown in blue; SaCysK + OAS shown in burgundy; EcCysK shown in yellow. The P(r) values are normalised by relative intensity for comparison of all enzymes on the same scale. Theoretical scattering shown in black in (A) and (C) and red in (E), calculated from OAS-free NgCysK crystal structure (9NLD). Guinier plots for all enzymes are inlaid in the scattering plots, enzymes without OAS are shown in blue, enzymes with OAS are shown in green and EcCysK is shown in black. A red trend line is present in all Guinier plots. The grey dashed lines in the Kratky plots represent the expected peak maxima for folded, globular proteins, where (qRg)^2^I(q)/I(0) = 1.104 in a q range of 0.05 – 0.1 Å^−1^ [85, 86].

**Table 7:**
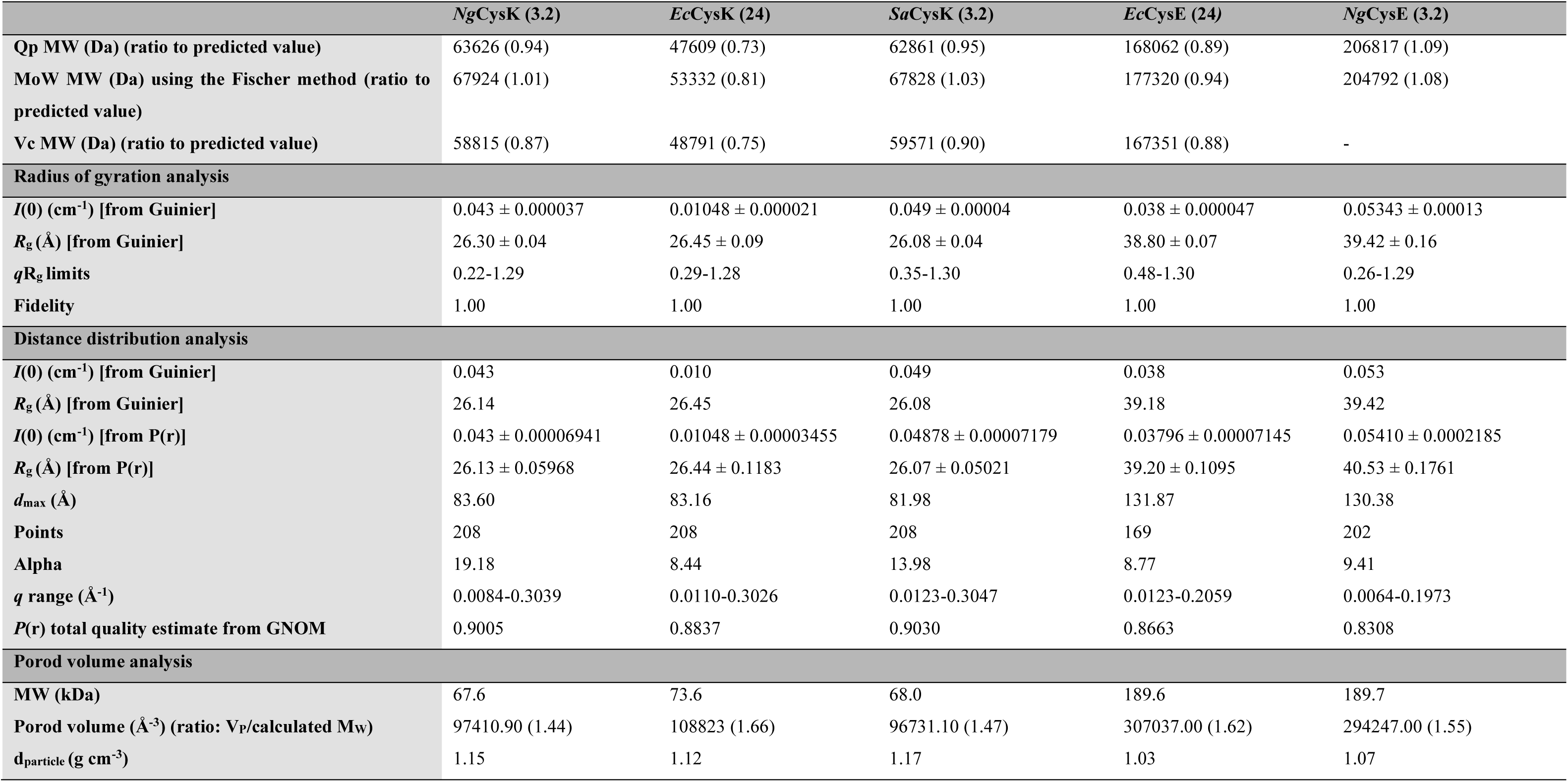
Structural parameters of Guinier fits, P(r) functions, MW estimates for *Ng*CysK*, Ng*CysE*, Ec*CysK*, Ec*CysE, and *Sa*CysK ran individually. 3.2 and 24 refer to the column volume used during SEX-SAXS run.

However, the MW calculations given by the Volume of correlation (V_c_), Fischer estimation (MoW), and the Porod invariant or dimensionless Kratky plot (Q_P_) methods, give more accurate values. The MW calculations for all three CysK enzymes give a ratio to the ProtParam calculated MW, consistent with that of a functional homodimer (EcCysK: 0.73-0.81 (Table 7); NgCsyK with and without OAS: 0.87-1.01 (Tables 7 and 8); SaCysK with and without OAS: ranging from 0.88-1.03 (Tables 7 and 8). The Guinier and P(r) calculated R_g_ values for all three enzymes are very similar, all ranging from 26.07-26.45 Å (Table 7), indicating the SaCysK and NgCysK behave in a similar manner to EcCysK, our “true” CysK, positive control. The EcCysK, NgCysK, and SaCysK SAXS profiles were fit with the calculated scattering from the NgCysK crystal structure (PDB ID: 9NLD). The theoretical scattering shows a poor fit with the scattering data collected for EcCysK (χ^2^ = 0.373), NgCysK (χ^2^ = 0.542), and SaCysK (χ^2^ = 0.616) (Supplementary Table 3), which is not unexpected due to the extremely flexible C-terminal tail of the CysK enzyme. However, in contrast, the theoretical scattering shows a good fit with the OAS bound NgCysK (χ^2^ = 0.977) and SaCysK (χ^2^ = 1.083) (Supplementary Table 3) scattering data. This indicates the binding of OAS into the active site may cause a conformation change to a more rigid conformation of the CysK enzyme, resulting in the scattering pattern more closely adhering to the theoretical scattering of the rigid NgCysK (9NLD) crystal structure. The scattering data of all three CysK enzymes, including the pairwise distribution analysis and R_g_ values from the Guinier analysis are consistent with literature SAXS data from EcCysK [83].

**Table 8.** Structural parameters of Guinier fits, P(r) functions, MW estimates for *Ng*CysK and *Sa*CysK in the presence of OAS.

|  | *NgCysK* + OAS | *SaCysK* + OAS |
| --- | --- | --- |
| **Qp MW (Da) (ratio to predicted value)** | 63093 (0.93) | 62861 (0.92) |
| **MoW MW (Da) using the Fischer method (ratio to predicted value)** | 60876 (0.90) | 67828 (0.99) |
| **Vc MW (Da) (ratio to predicted value)** | 61531 (0.91) | 59571 (0.88) |
| **Radius of gyration analysis** |  |  |
| *I*(0) (cm<sup>-1</sup>) | 0.025 ± 0.0001 | 0.031 ± 0.0001 |
| *R<sub>g</sub>* (Å) | 26.11 ± 0.18 | 25.96 ± 0.16 |
| *qR<sub>g</sub>* limits (Å<sup>-1</sup>) | 0.22-1.29 | 0.28-1.25 |
| Fidelity | 0.80 | 0.82 |
| **Distance distribution analysis** |  |  |
| *I*(0) (cm<sup>-1</sup>) [from Guinier] | 0.025 | 0.031 |
| *R<sub>g</sub>* (Å) [from Guinier] | 25.90 | 26.11 |
| *I*(0) (cm<sup>-1</sup>) [from P(r)] | 0.02520 ± 0.0001026 | 0.03098 ± 0.00008989 |
| *R<sub>g</sub>* (Å) [from P(r)] | 25.89 ± 0.1487 | 26.11 ± 0.07990 |
| *d<sub>max</sub>* (Å) | 77.69 | 81.98 |
| Points | 170 | 169 |
| Alpha | 0.1294 | 12.12 |
| *q* range (Å<sup>-1</sup>) | 0.0083-0.3064 | 0.0083-0.3084 |
| *P*(r) total quality estimate from GNOM | 0.98 | 0.98 |
| **Porod volume analysis** |  |  |
| MW (kDa) | 67.58 | 68.02 |
| Porod volume (Å<sup>-3</sup>) (ratio: *V<sub>p</sub>*/calculated MW) | 95074.90 (1.41) | 94602.70 (1.39) |
| *d<sub>particle</sub>* (g cm<sup>-3</sup>) | 1.18 | 1.19 |

Comparison of the NgCysK and SaCysK SAXS profiles in the presence and absence of OAS show negligible differences (Figure 5), indicating no major change in conformation or flexibility of the enzyme [68]. This indicates there is no change in the compactness of the enzymes when OAS is bound, as shown by the R_g_ values derived from Guinier analysis. Values from the Guinier analysis tool (radius of gyration analysis) are used as this assumes a globular shape [84] of which NgCysK and SaCysK are. There is no change in R_g_, from holo NgCysK to NgCysK with OAS bound (26.30 ± 0.04 - 26.11 ± 0.18) or from holo SaCysK to SaCysK with OAS bound (26.08 ± 0.04 - 25.96 ± 0.16) (Tables 7 and 8).

The Kratky plots for both NgCysK and SaCysK with and without OAS, alongside EcCysK indicate a bell-shaped curve, consistent with the expected shape exhibited by a typical folded protein (Figure 5). This confirms that NgCysK and SaCysK enzymes remain folded in the presence of OAS.

### Formation of the Cysteine Synthase Complex (CSC)

There are three main mechanisms for regulating L-cysteine metabolism and therefore sulfur flux. The first is transcriptional regulation by CysB, and the second is feedback inhibition of CysE by L-cysteine which is seen in other CysE homologues [87, 88], including *N. gonorrhoeae* [53]. The final, which we will investigate here, is formation of the CSC which occurs within many bacterial and plant species [26, 31, 43–45, 51] and is hypothesised to modulate sulfur flux [39, 89, 90]. Due to the use of the CSC in modulating sulfur flux, combined with both, L-cysteine mediated CysE inhibition and a lack of the sulfate reduction pathway, it is of particular interest whether *N. gonorrhoeae* and *S. aureus* can form the CSC. Using Clustal Omega [91], a sequence alignment of CysE bacterial homologues was generated and analysed using ESPript 3.0 (Figure 6) [92]. This analysis showed the CysE from *N. gonorrhoeae* (*Ng*CysE) and *S. aureus* (*Sa*CysE), had moderate (49.51%) and low to moderate (38.69%) sequence similarities to other CysE homologues, respectively (Figure 6). The C-terminus four peptide fragment, GDGI was well conserved across three species with DFMI and DYII being the C-terminal tetrapeptide sequence in NgCysE and *Sa*CysE, respectively. The conservation of the C-terminal isoleucine in both species indicates their potential for CSC formation. The CysK active site has also been confirmed as the anchor for CSC formation [4]. The CSC is an important regulator of the cysteine biosynthesis pathway and has yet to be characterised in *N. gonorrhoeae* or *S. aureus*.

**Figure 6:**
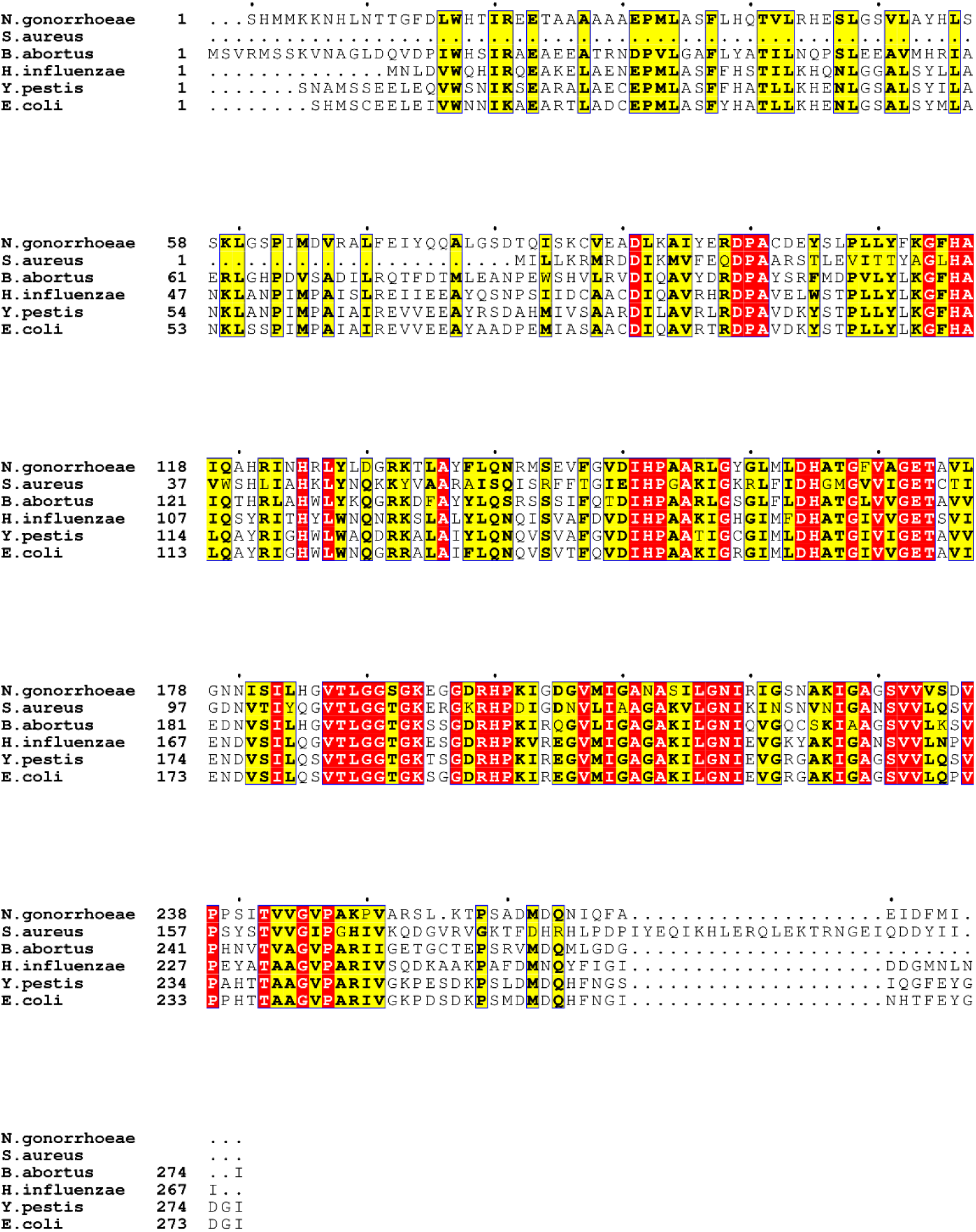
CysE protein sequence alignment. The *N. gonorrhoeae* and *S. aureus* CysE amino acid sequences were aligned to the following CysE homologues; *Brucella abortus* (*Ng*CysE 47.96% and *Sa*CysE 41.27% sequence similarity); *Haemophilus influenzae* (*Ng*CysE 51.89% and *Sa*CysE 39.49% sequence similarity); *Yersinia pestis* (*Ng*CysE 53.14% and *Sa*CysE 36.92% sequence similarity); *Escherichia coli* (*Ng*CysE 53.70% and *Sa*CysE 36.92% sequence similarity); and *Ng*CysE and *Sa*CysE have 39.18% sequence similarity to each other. Highly conserved residues are highlighted red, whilst less conserved residues are highlighted yellow. Sequence alignment created using Clustal Omega [91]. Figure created using ESPript 3.0 [92].

Characterising the CSC would provide further insight into elucidating the sulfur assimilation and synthesis of L-cysteine in *N. gonorrhoeae* and *S. aureus*. However, we were unable to purify the SaCysE enzyme, therefore, only the NgCSC formation was tested using *E. coli* as a positive control as it is known to form the CSC [31, 54].

### Investigation of CSC formation by gel filtration chromatography

Formation of the CSC was investigated by gel filtration chromatography using freshly purified CysK and CysE. Both proteins were purified individually, and analysed using a calibrated gel filtration column. NgCysE elutes from an Enrich 650 analytical gel filtration column (BioRad) at 12.28 ml, corresponding to a molecular weight of 193.8 kDa, consistent with an NgCysE hexamer (Figure 7A). NgCysK elutes at 14.6 ml, corresponding to a molecular weight of 52.862 kDa, consistent with a dimer (Figure 7C). EcCysE elutes from an Enrich 650 analytical gel filtration column (BioRad) at 13.4 ml, corresponding to a molecular weight of 198.5 kDa, consistent with an EcCysE hexamer (Figure 7B). EcCysK elutes at 15 ml, corresponding to a molecular weight of 80.1 kDa, consistent with a dimer (Figure 7D).

**Figure 7:**
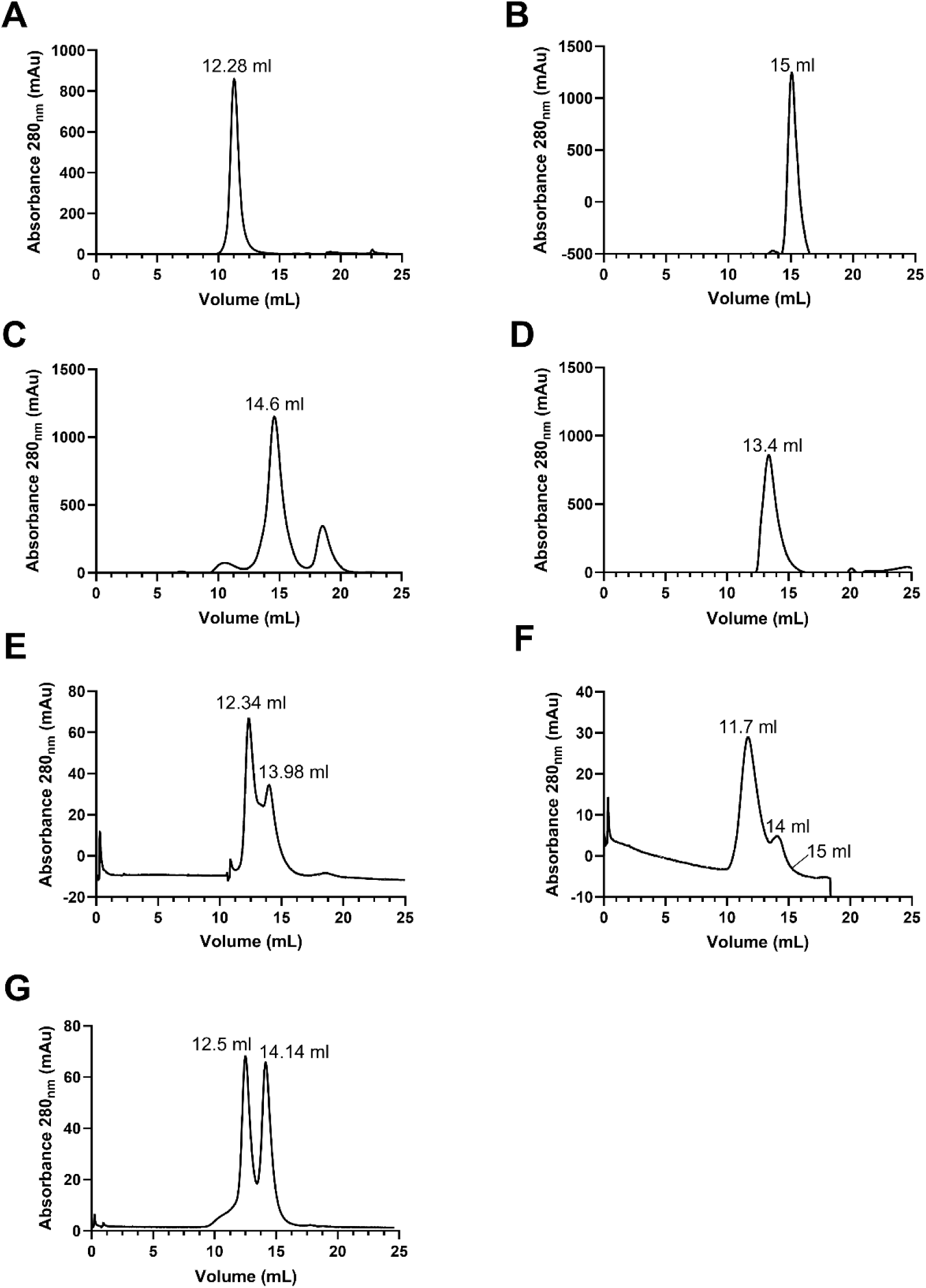
Monitoring formation of the CSC in *N. gonorrhoeae* and *E. coli* by gel filtration chromatography

NgCysE and NgCsyK were mixed and analysed by gel filtration chromatography at 3:2 and 1:1 monomeric molar ratios. The homologous CSC from *E. coli* indicates the composition to be one CysE hexamer and two CysK dimers (3:2 monomeric molar ratio) [31]. Given this conformation, the NgCSC formation would give an elution peak of ∼332 kDa corresponding to an elution volume of 11.9 ml, prior to the NgCysE and NgCysK elution peaks seen, however, we observe no such peak indicating no CSC formation (Figure 7).

In comparison, given the established 3:2 ratio of CysE to CysK in *E. coli* [31], the EcCSC formation would give an elution peak of ∼337 kDa corresponding to an elution volume of 11.8 ml, prior to the EcCysE and EcCysK elution peaks seen. Given the presence of this peak in the elution profile (Figure 7F), this indicates successful formation of the EcCSC.

### Investigation of CSC formation by SAXS

Following gel filtration chromatography experiments, we looked at the potential CSC formation in *N. gonorrhoeae* in comparison to our *E. coli* positive control using SEC-SAXS. Both the EcCSC and NgCSC forming enzymes were run as per Table 2. Visual inspection of the UV trace of the SEC run prior to beam exposure and the CHROMIXS scattering profile, shows the presence of three distinct peaks in the EcCSC run compared to the presence of only two distinct peaks in the NgCSC run (Figure 8). The first peak seen in the EcCSC profiles indicates the expected larger CSC followed by the elution of the excess (hasn’t formed the CSC) EcCysE, and the third and final peak being the excess EcCysK (Figure 8). This was expected as the CSC formation functions in an equilibrium [50, 93], unless driven by other factors such as the presence of OAS, which can dissociate the complex at concentrations upwards of 50 µM [43, 50, 79] or the presence of sulfide, which promotes complex formation [39]. In comparison, visual inspection of the UV trace and CHROMIXS scattering for the NgCSC run, shows only the elution peaks for NgCysE and NgCysK (Figure 8A).

**Figure 8:**
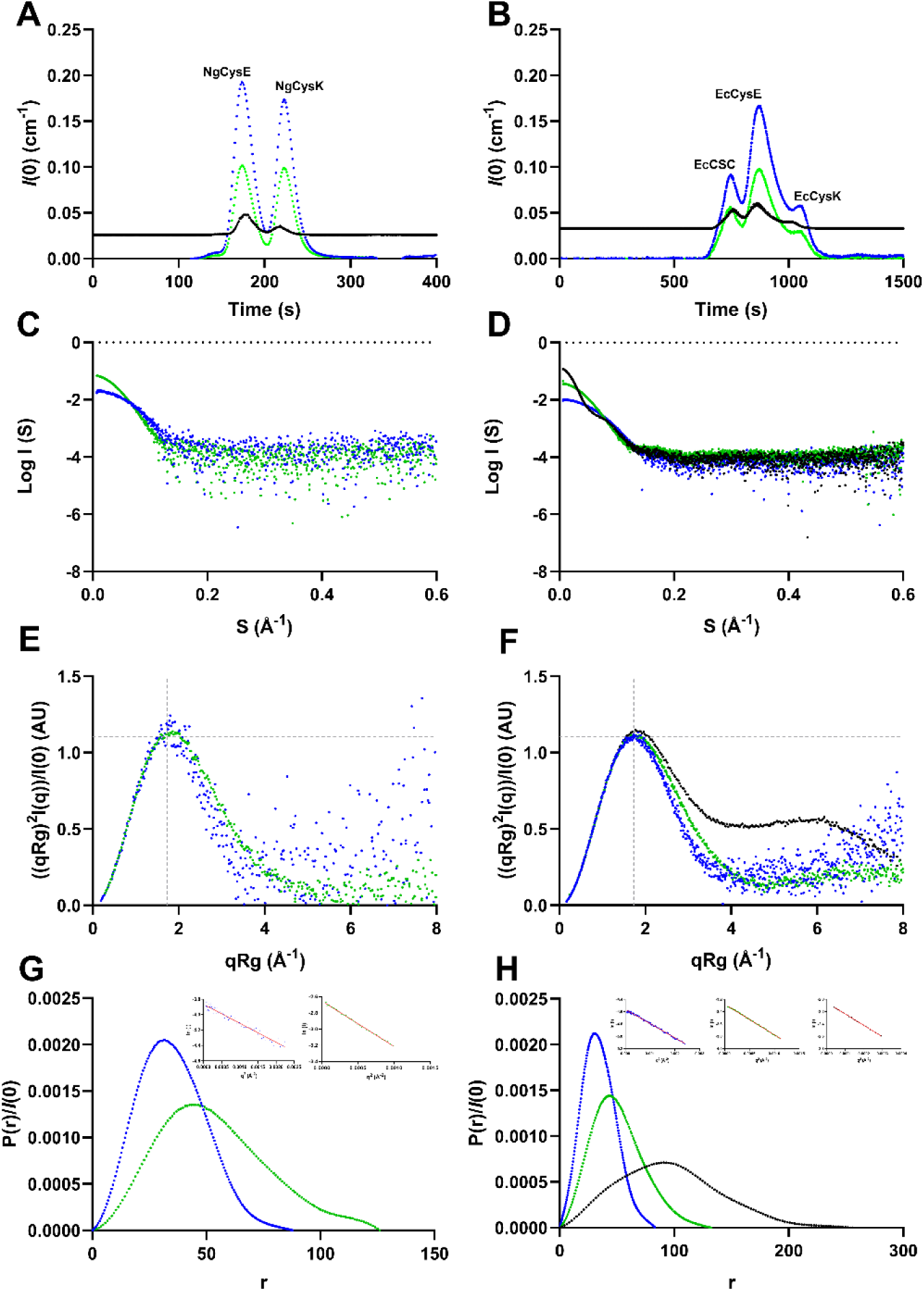
SAXS profiles of attempted CSC formation in *N. gonorrhoeae* and *E. coli.* (A) UV trace and CHROMIXS scattering of attempted NgCSC formation. (B) UV trace and CHROMIXS scattering of EcCSC formation. (C) Scattering profiles of NgCysK (blue circles) and NgCysE (green circles) from attempted NgCSC formation. (D) Scattering profiles of of EcCysK (blue circles), EcCysE (green circles), and EcCSC (black circles) from EcCSC formation. (E) Kratky plot of NgCysK (blue circles) and NgCysE (green circles) from attempted NgCSC formation. (F) Kratky plot of EcCysK (blue circles), EcCysE (green circles), and EcCSC (black circles) from EcCSC formation. (G) Pair distance distribution function (P(r)) analysis of NgCysK (blue circles) and NgCysE (green circles) from attempted NgCSC formation. (H) Pair distance distribution function (P(r)) analysis of EcCysK (blue circles), EcCysE (green circles), and EcCSC (black circles) from EcCSC formation. Individual enzymes present in the UV trace and CHROMIXS scattering are labelled in black. UV A_280_ trace is shown in blue. UV A_260_ trace is shown in green. CHROMIXS scattering is shown in black. The grey dashed lines in the Kratky plots represent the expected peak maxima for folded, globular proteins, where (qRg)^2^I(q)/I(0) = 1.104 in a q range of 0.05 – 0.1 Å^−1^ [85, 86]. Guinier plots for all enzymes are inlaid on the P(r) distributions. A red trend line is present in all Guinier plots.

Analysis of the scattering data from these runs also shows a distinct difference between the EcCSC and NgCSC runs (Figure 8 and Table 9). The scattering points from each of the peaks were isolated and analysed separately, and once extrapolated, showed a distinctly different scattering pattern for the EcCSC in comparison to the individual enzymes (EcCysE and EcCysK) scattering patterns (Figure 8D). The scattering of the individual enzymes is typical of a folded globular protein, further consolidated by the bell shape curve of their Kratky plots (Figure 8F), and the P(r) distribution analysis (Figure 3.8H). However, the EcCSC peak has a distinct scattering pattern indicating the presence of a large molecule with multiple species (i.e. EcCysE and EcCysK) together in a complex (Figure 8D). This is also evident in the EcCSC Kratky plot and P(r) distribution analysis (Figure 8).

**Table 9:** Structural parameters of Guinier fits, P(r) functions, MW estimates for CSC formation.

|  | *NgCysE* (3.2 ml column) | *NgCysK* (3.2 ml column) | *EcCSC* (24 ml column) |
| --- | --- | --- | --- |
| **Qp MW (Da) (ratio to predicted value)** | 216494 (1.14) | 64574 (0.96) | 742926 (2.21) |
| **MoW MW (Da) using the Fischer method (ratio to predicted value)** | 190215 (1.00) | 66520 (0.98) | - |
| **Vc MW (Da) (ratio to predicted value)** | 208954 (1.10) | 60934 (0.90) | - |
| **Radius of gyration analysis** |  |  |  |
| ***I*(0) (cm<sup>-1</sup>)** | 0.07 ± 0.0002 | 0.021 ± 0.00018 | 0.1284 ± 0.00017 |
| ***R<sub>g</sub>* (Å)** | 40.94 ± 0.19 | 27.09 ± 0.39 | 74.85 ± 0.14 |
| ***qR<sub>g</sub>* limits (Å<sup>-1</sup>)** | 0.34-1.26 | 0.21-1.30 | 0.53-1.30 |
| **Fidelity** | 0.75 | 0.72 | 0.99 |
| **Distance distribution analysis** |  |  |  |
| ***I*(0) (cm<sup>-1</sup>) [from Guinier]** | 0.070 | 0.021 | 0.128 |
| ***R<sub>g</sub>* (Å) [from Guinier]** | 41.15 | 26.87 | 74.85 |
| ***I*(0) (cm<sup>-1</sup>) [from P(r)]** | 0.07022 ± 0.0001692 | 0.02085 ± 0.0001642 | 0.1284 ± 0.0002387 |
| ***R<sub>g</sub>* (Å) [from P(r)]** | 41.16 ± 0.1118 | 26.87 ± 0.2545 | 74.91 ± 0.1793 |
| ***d<sub>max</sub>* (Å)** | 144.74 | 81.91 | 255.85 |
| **Points** | 134 | 167 | 122 |
| **Alpha** | 14.60 | 3.31 | 18.69 |
| ***q* range (Å<sup>-1</sup>)** | 0.0083-0.1947 | 0.0077-0.2952 | 0.0071-0.1074 |
| ***P*(r) total quality estimate from GNOM** | 0.8094 | 0.9257 | 0.9009 |
| **Porod volume analysis** |  |  |  |
| **MW (kDa)** | 189.7 | 67.6 | 336.72 |
| **Porod volume (Å<sup>-3</sup>) (ratio: V<sub>P</sub>/calculated MW)** | 313606.00 (1.65) | 99147.40 (1.47) | 1465780.00 (4.35) |
| ***d<sub>particle</sub>* (g cm<sup>-3</sup>)** | 1.00 | 1.13 | 0.38 |

Given the conservation of the C-terminal isoleucine in *N. gonorrhoeae* and the confirmation of their successful folding into functional globular proteins, the potential for CSC formation is there. However, through both gel filtration chromatography and SAXS analysis, we have confirmed the inability of *N. gonorrhoeae* to form the CSC *in vitro*. It is therefore possible that the CSC is completely unable to form in *N. gonorrhoeae*. This would result in a CysK enzyme that is not inhibited by CSC formation, allowing an increase in L-cysteine production. Additionally, due to the unregulated OAS consumption by CysK, sulfate acquisition genes would be downregulated. Given the necessity of reducing compounds such as glutathione [18], therefore creating high demand for L-cysteine, it may be advantageous for *N. gonorrhoeae* to be incapable of CSC formation. This lack of CSC formation may in fact be linked to the inability of *N. gonorrhoeae* to reduce sulfate to sulfide, and that the transcriptional regulator CysB controls two of the deleted genes from this pathway via operon control.

## Conclusion

Sulfur metabolism across plant and bacterial species is vital to many life cycles within our planetary ecosystem. Investigating the various avenues of sulfur metabolism will not only garner a profoundly deeper understanding of the planet we live on, but offer the capability to develop new medicines, new environmentally friendly production methods, new engineering processes, or even ways to break down and remediate already existing environmental damage. Given that L-cysteine biosynthesis is at the center of sulfur metabolism, particularly in bacterial organisms, investigating bacterial sulfur metabolism, particularly of two pathogens with the same non-functional sulfur reduction pathway such as *N. gonorrhoeae* and *S. aureus* is of vital importance.

Here we have presented the kinetic and SEC-SAXS analysis of CysK from both *N. gonorrhoeae* and *S. aureus*. Small angle X-ray scattering analysis confirms that both enzymes function as dimers in solution with no other oligomeric states detected. We have characterised the kinetics of both enzymes, and hypothesise a bi-bi ping-pong mechanism, with *O-*acetylserine being first to bind, and once the acetyl group is released, sulfide enters the active site replacing the acetyl group with a thiol group, to form L-cysteine. We suggest positive cooperativity at low concentrations of OAS with a plateau beginning at concentrations ≥5 mM, and positive cooperativity at Na_2_S concentrations of ≤15 mM for NgCysK. We suggest substrate inhibition at OAS concentrations ≥5 mM and Na_2_S concentrations ≥12 mM for SaCysK. Both enzymes have a higher affinity for OAS (NgCysK: 1.541 mM, SaCysK: 1.378mM) compared to Na_2_S (NgCysK: 23.620 mM, SaCysK: 2.306 mM), which is intriguingly similar to the plant *O-*acetylserine sulfhydrylase isozyme A and C from *D. innoxia* [72], but not to other bacterial homologues [34, 71, 75]. The kinetic relationship to both substrates is substantially varied across CysK homologues, however, some similarities do exist. NgCysK exhibits positive cooperativity for both OAS (also seen in *D. innoxia* [72], and Na_2_S (also seen in *P. vulgaris* and *D. innoxia* [71, 72]). Conversely, SaCysK exhibits substrate inhibition for both OAS (also seen in *D. innoxia* and *S.* typhimurium [34, 72, 75]) and Na_2_S (also seen in *P. vulgaris, D. innoxia* OASS (isozyme C), *P. polyanthus,* and *S. typhimurium* [71, 72, 75]). The cooperativity seen in NgCysK indicates an interconnectedness between active sites of the NgCysK dimer. This is not observed in the SaCysK. SaCysK and NgCysK are both incapable of utilising thiosulfate as a sulfur donor in L-cysteine biosynthesis, confirming their characterisation as CysK enzymes.

Binding of OAS into the active sites of both enzymes causes some manner of conformational shift or rigidity as indicated by the improved chi-square value of both NgCysK and SaCysK in the presence of 10 mM OAS, compared to without OAS. However, this substrate binding does not cause a change in the overall flexibility or density of the enzymes, indicating a small internal conformation change.

Further elucidation of the mechanisms of sulfur metabolism also led us to the investigation of CSC formation. Using both, gel chromatography and SAXS analysis, we have shown that *N. gonorrhoeae* is incapable of CSC formation *in vitro.* This raises further questions as to the regulation of sulfur metabolism in this organism. The lack of CSC formation would mean a lack of NgCysK inhibition which may be an evolutionary advantage for *N. gonorrhoeae,* due to the high demand for L-cysteine production to produce vital reducing compounds such as glutathione. Combined with the L-cysteine feedback inhibition of NgCysE [53], this may indicate an alternate mechanism of sulfur flux regulation in *N. gonorrhoeae*.

In bacteria, all primary pathways for inorganic sulfur assimilation converge on L-cysteine biosynthesis. Our two organisms, *S. aureus* and *N. gonorrhoeae,* both lack the ability to use inorganic sulfate as a sulfur source, but can fulfil their requirement for sulfur with thiosulfate [12, 13]. Both CysK enzymes from these pathogenic bacteria are unable to use thiosulfate for L-cysteine production, indicating the presence of an alternate sulfur acquisition pathway in both *S. aureus* and *N. gonorrhoeae.* Given the inability of *N. gonorrhoeae* to form the CSC, combined with an inability to reduce sulfate or use thiosulfate for L-cysteine production, there are many more questions as to the mechanisms of sulfur regulation, acquisition and metabolism in *N. gonorrhoeae*.

Our NgCysK and SaCysK kinetics, alongside the SAXS analysis of both enzymes gives unique insight into the function of two bacteria lacking the same sulfate reduction pathway. This, combined with the inability of *N. gonorrhoeae* to form the CSC, indicates another yet undetermined sulfur metabolising pathway that will lead to a deeper understanding of our ecosystem, and may lead to advances in medicinal, climate change, and biotechnological research.

## Notes

### Competing Interest Statement

The authors have declared no competing interest.

