## Supplementary Data for "Characterisation of O-acetylserine sulfhyrdrylase (CysK) enzymes from bacteria lacking a sulfate reduction pathway"

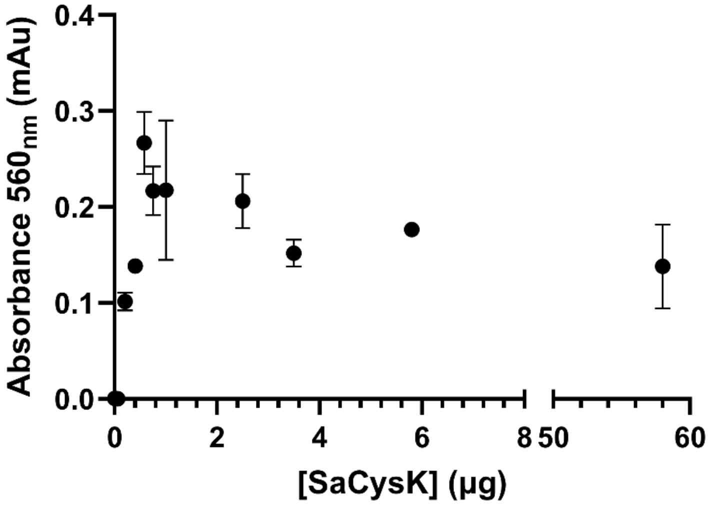


Supplementary Figure 1: Absorbance readings of Cysteine produced from various SaCysK concentrations. Given the error in the points above 0.2 μg, and the plateau at ~0.15 mAu, 0.2 μg was selected as the appropriate concentration for further testing.

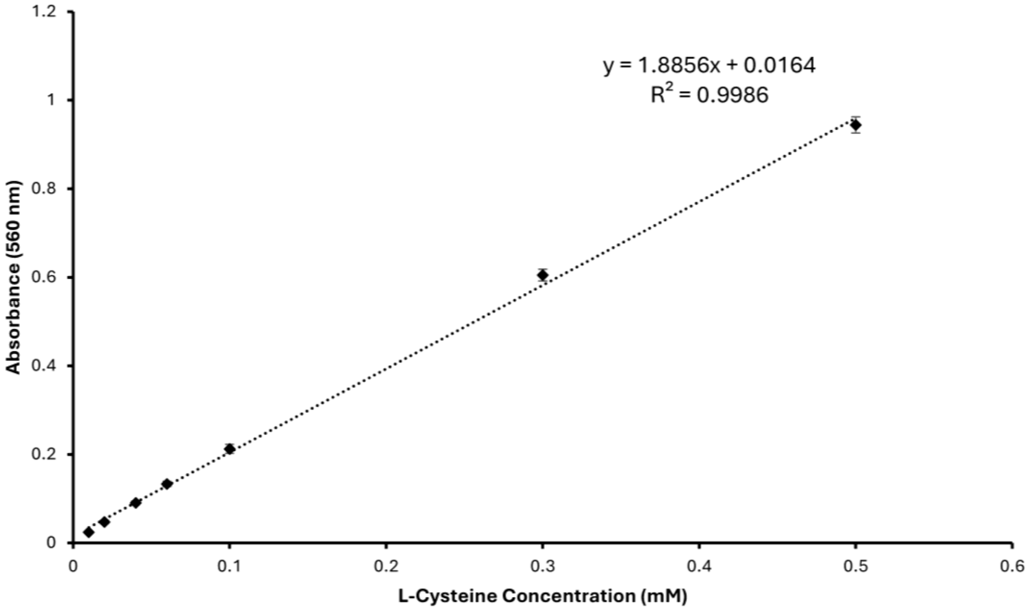


Supplementary Figure 2: Standard curve of L-cysteine. Concentration (mM) of L-cysteine plotted on the x-axis. Absorbance ($A_{560}$) of L-cysteine plotted on the y-axis. Cysteine produced in kinetic assays was calculated from absorbance readings ($A_{560}$) using the equation from the line of best fit.

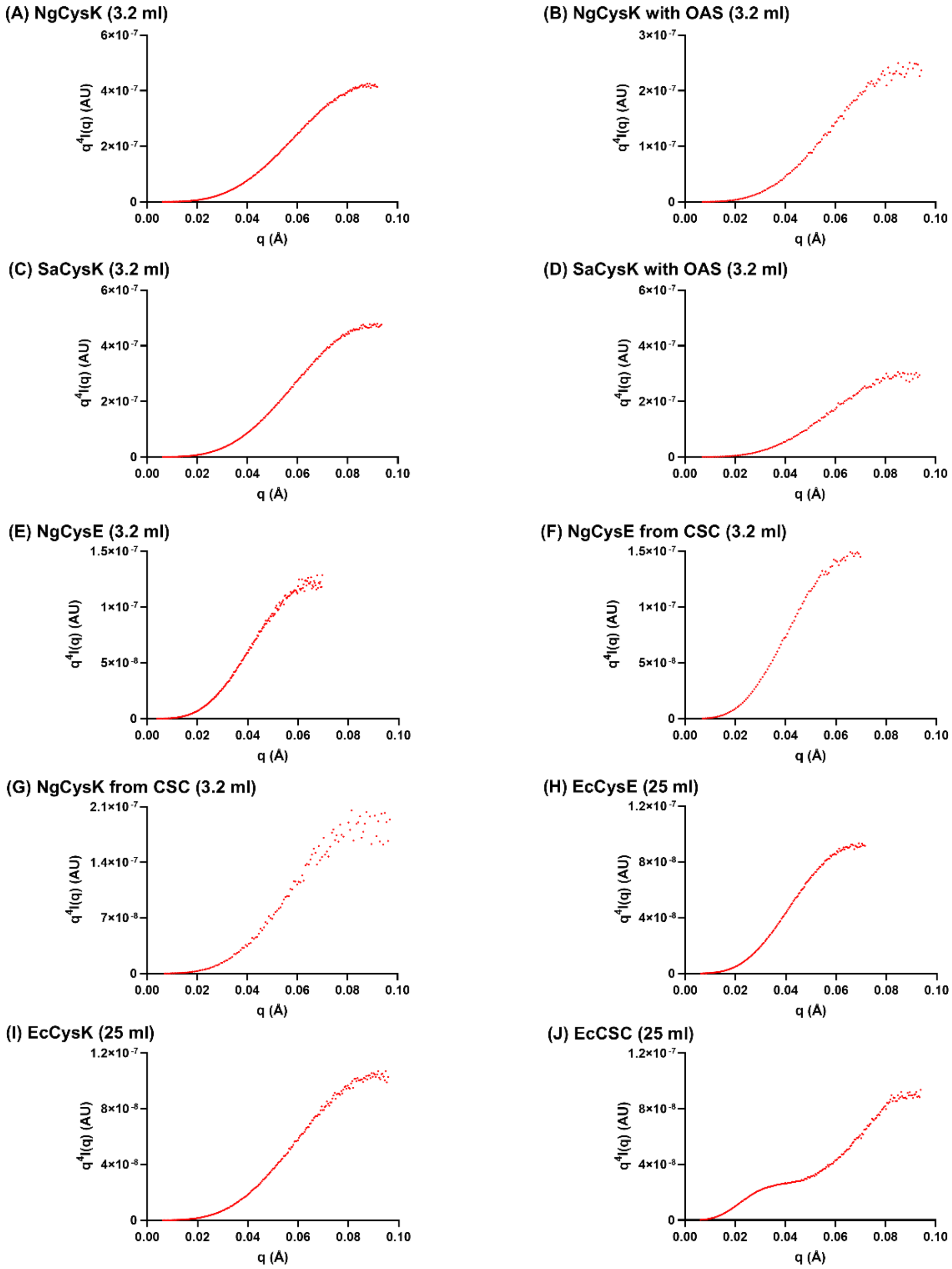


Supplementary Figure 3: Porod-debye plots defining the q range used for determining the Porod volume ($V_p$)

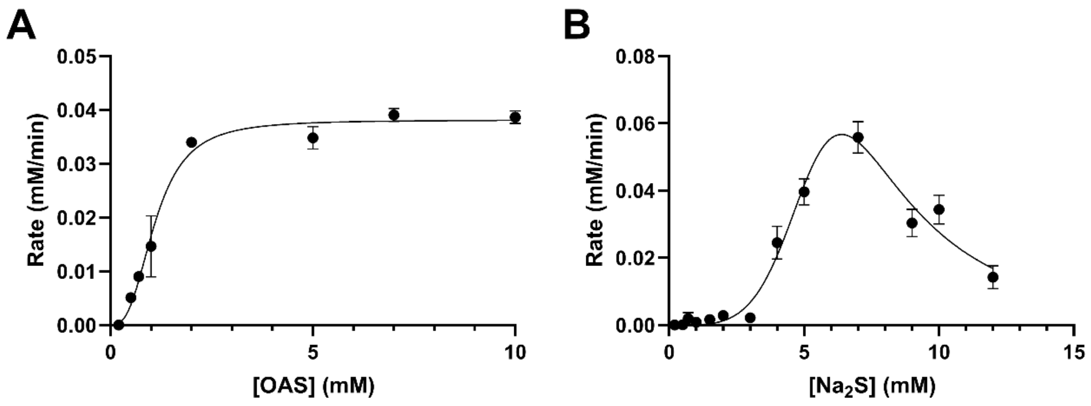


Supplementary Figure 4: Kinetic analysis of NgCysK substrates OAS and $Na_2S$ in the presence of glycerol. (A) Allosteric sigmoidal fit (occurs when the enzyme has cooperative subunits) for OAS. (B) Constrained allosteric sigmoidal substrate inhibition fit (occurs when the enzyme has cooperative subunits and exhibits cooperativity at low substrate concentrations and inhibition at high substrate concentrations) for $Na_2S$. OAS and $Na_2S$ assays in the presence of glycerol were collected at saturating concentrations of 10 mM OAS and 7 mM $Na_2S$, respectively. Plotted data points represent mean alongside SEM of three replicates.

Supplementary Table 1: Kinetic parameters of NgCysK in the presence of glycerol

| Parameter | *O*-acetylserine (OAS)[a] | Sodium sulfide ($Na_2S$)[a] |
|---|---|---|
| $V_{max}$ (mM.min$^{-1}$) | 0.038 ± 0.002 | 2.530 ± 4.601 |
| $K_M$ (mM) | - | 12.43 ± 8.133 |
| $K_{half}$ (mM) | 1.101 ± 0.138 | - |
| $k_{cat}$ (s$^{-1}$) [b] | (2.148 ± 0.130) x10$^5$ | (1.425 ± 2.590) x 10$^7$ |
| $k_{cat}/K_M$ (M.s$^{-1}$) | (1.951 ± 0.272) x 10$^8$ | (1.146 ± 2.217) x 10$^9$ |
| $K_i$ (mM) | - | 5.146 ± 5.012 |
| h | 2.810 ± 0.899 | 4.159 ± 1.549 |
| $R^2$ | 0.9795 | 0.9573 |

[a] Error is SEM of three replicates

[b] $k_{cat}$ calculated by dividing the rate (M.s$^{-1}$) by enzyme concentration using the concentration of the NgCysK dimer (33.791 kDa)

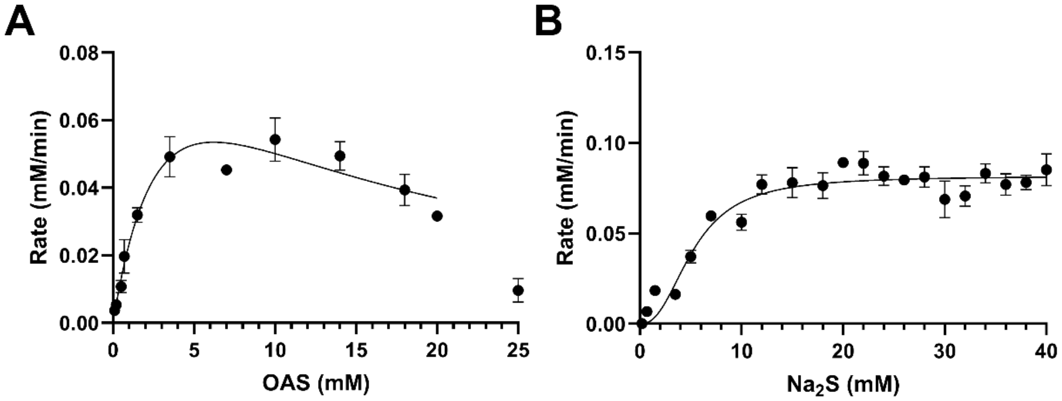


Supplementary Figure 5: Kinetic analysis of SaCysK substrates OAS and $Na_2S$ in the presence of glycerol. (A) Substrate inhibition fit (occurs when the enzyme exhibits inhibition at high substrate concentrations) for OAS. (B) Allosteric sigmoidal fit (occurs when the enzyme has cooperative subunits) for $Na_2S$. OAS and $Na_2S$ assays in the presence of glycerol were collected at saturating concentrations of 10 mM OAS and 15 mM $Na_2S$, respectively. Plotted data points represent mean alongside SEM of three replicates.

Supplementary Table 2: Kinetic parameters of SaCysK in the presence of glycerol

| Parameter | *O*-acetylserine (OAS)[a] | Sodium sulfide ($Na_2S$)[a] |
|---|---|---|
| $V_{max}$ ($mM.min^{-1}$) | 0.129 ± 0.087 | 0.082 ± 0.004 |
| $K_M$ (mM) | 4.387 ± 4.404 | - |
| $K_{half}$ (mM) | - | 5.266 ± 0.679 |
| $k_{cat}$ ($s^{-1}$) [b] | $(7.313 \pm 4.952)$ x$10^5$ | $(4.649 \pm 0.024)$ x $10^5$ |
| $k_{cat}/K_M$ ($M.s^{-1}$) | $(1.667 \pm 2.019)$ x $10^8$ | $(8.830 \pm 1.226)$ x $10^7$ |
| $K_i$ (mM) | 8.810 ± 7.348 | - |
| h | - | 2.397 ± 0.906 |
| $R^2$ | 0.9347 | 0.9184 |

[a] Error is SEM of three replicates
[b] $k_{cat}$ calculated by dividing the rate ($M.s^{-1}$) by enzyme concentration using the concentration of the SaCysK dimer (34.011 kDa)

Supplementary Table 3: Crysol results and parameters. 3.2 and 24 refer to the column volume used during SEC-SAXS run.

| | ***Ng*CysK (3.2)** | ***Ec*CysK (24)** | ***Sa*CysK (3.2)** | ***Ec*CysE (24)** | ***Ng*CysE (3.2)** | ***Ng*CysK + OAS (3.2)** | ***Sa*CysK + OAS (3.2)** |
|---|---|---|---|---|---|---|---|
| **Symmetry, anisotropy assumptions** | P2 | P2 | P2 | P6 | P6 | P2 | P2 |
| **Red. $\chi^2$, CorMap, Anderson-darling, Probability of fit** | 0.542, 75, 24.986, 1.000 | 0.373, 22, 52.319, 1.000 | 0.616, 89, 14.997, 1.000 | 2.513, 124, 45.384, 0.0 | 0.542, 99, 37.826, 1.0 | 0.977, 15, 0.918, 0.654 | 1.083, 20, 3.126, 0.046 |
| **Optimal hydration shell contrast (Dro, e Å$^{-3}$)** | 0.018 | 0.014 | 0.021 | 0.100 | 0.065 | 0.022 | 0.025 |
| **Predicted $R_g$ (Å)** | 24.65 | 24.65 | 24.65 | 34.43 | 34.71 | 24.65 | 24.65 |
| **Optimal Excluded volume (Vol, Å$^3$)** | 70813 | 70813 | 70813 | 211505 | 195566 | 70813 | 70813 |
| **Crysol (default parameters used)** | | | | | | | |
| **Maximum angle** | 0.5 Å$^{-1}$ | 0.5 Å$^{-1}$ | 0.5 Å$^{-1}$ | 0.5 Å$^{-1}$ | 0.5 Å$^{-1}$ | 0.5 Å$^{-1}$ | 0.5 Å$^{-1}$ |
| **Number of points** | 501 | 501 | 501 | 501 | 501 | 501 | 501 |
| **Solvent density** | 0.334 e/Å$^3$ | 0.334 e/Å$^3$ | 0.334 e/Å$^3$ | 0.334 e/Å$^3$ | 0.334 e/Å$^3$ | 0.334 e/Å$^3$ | 0.334 e/Å$^3$ |
| **Hydration shell constant** | 0.03 e/Å$^3$ | 0.03 e/Å$^3$ | 0.03 e/Å$^3$ | 0.03 e/Å$^3$ | 0.03 e/Å$^3$ | 0.03 e/Å$^3$ | 0.03 e/Å$^3$ |
| **Maximum order of harmonics** | 70 | 70 | 70 | 70 | 70 | 70 | 70 |
| **Order of Fibonacci grid** | 17 | 17 | 17 | 17 | 17 | 17 | 17 |
| **Shell type** | Directional | Directional | Directional | Directional | Directional | Directional | Directional |

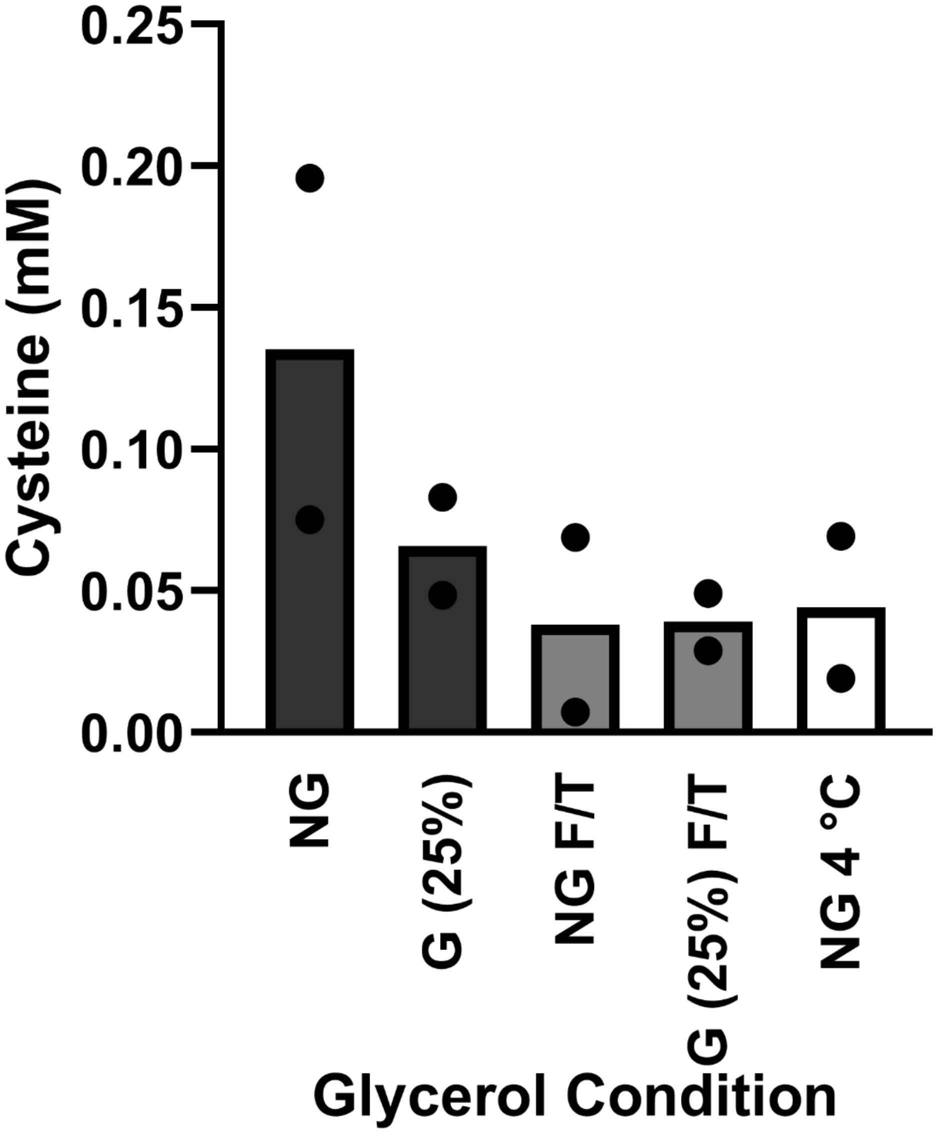


Supplementary Figure 5: Column graph displaying the effect of glycerol on enzyme activity under varying conditions at 0.704 and 1.408 ug. NG: fresh NgCysK without glycerol. G (25%): fresh NgCysK with 25% (v/v) glycerol. NG F/T: freeze/thawed NgCysK without glycerol. G (25%) F/T: freeze/thawed NgCysK with 25% (v/v) glycerol. NG 4 ºC: NgCysK without glycerol refrigerated for 3 days. Each concentration was collected in duplicate under each condition. Points shown under each condition are the mean of each enzyme concentration. Columns shown are the mean of both concentrations in that condition.

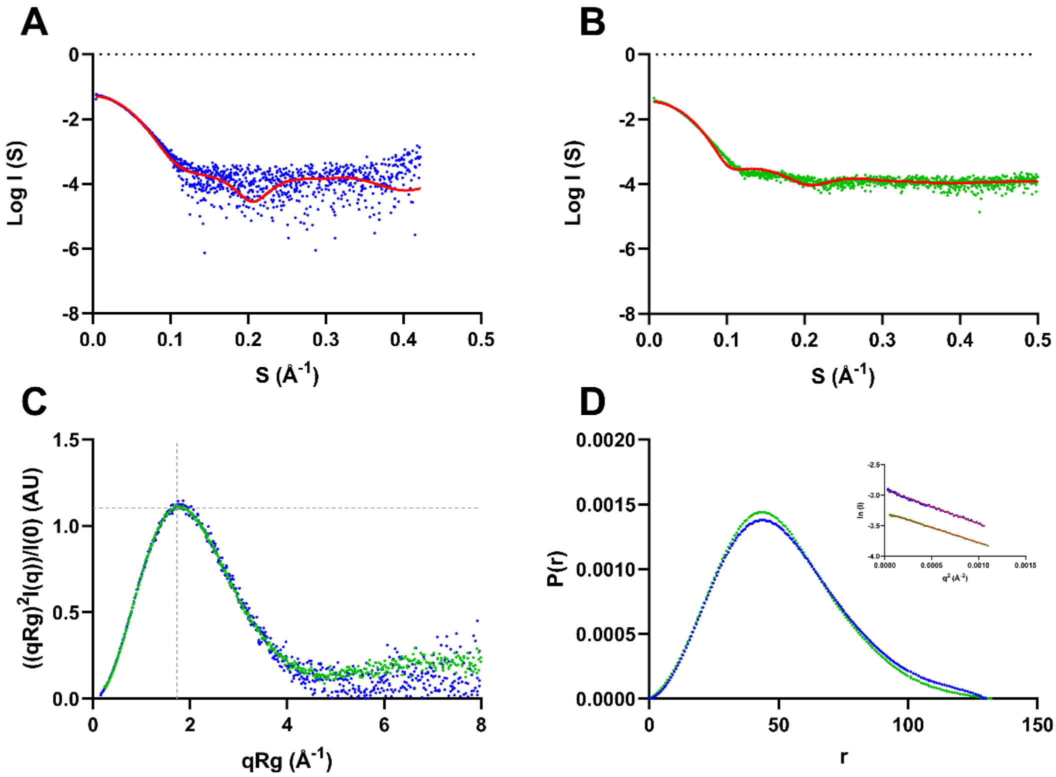


Supplementary Figure 1: SAXS profiles of NgCysE (all blue curves) and EcCysE (all green curves). (A) Scattering plot of NgCysE with theoretical scattering fit of NgCysE (6WYE) shown in red. (B) Scattering plot of EcCysE with theoretical scattering fit of EcCysE (1T3D) shown in red. (C) Kratky plots of NgCysE (blue) and EcCysE (green). The grey dashed lines represent the expected peak maxima for folded, globular proteins, where $(qRg)^2I(q)/I(0) = 1.104$ in a q range of 0.05 – 0.1 Å$^{-1}$ [85, 86]. (D) Pair distance distribution function (P(r)) analysis of NgCysE (blue) and EcCysE (green) with inlaid Guinier plots. A red trend line is present in both Guinier plots. The P(r) values are normalised by relative intensity for comparison of both enzymes on the same scale.

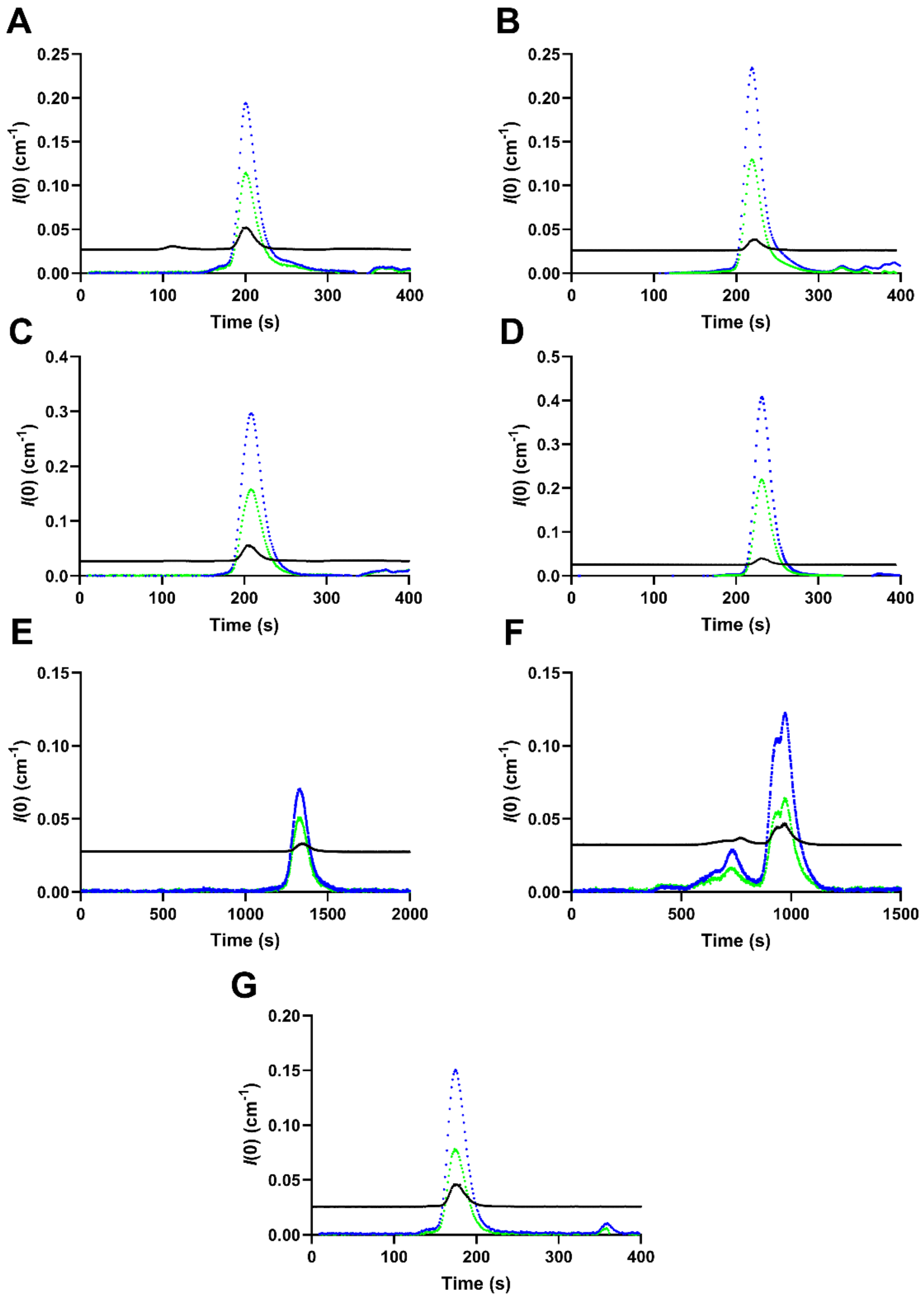


Supplementary Figure 2: UV traces and CHROMIXS scattering of all individual enzymes tested, excluding those involved in CSC formation. (A) NgCysK. (B) NgCysK + OAS. (C) SaCysK. (D) SaCysK + OAS. (E) EcCysK. (F) EcCysE. (G) NgCysE. For all curves, UV $A_{280}$ trace is shown in blue, UV $A_{260}$ is shown in green, and CHROMIXS scattering is shown in black.
